# Bridging themes: short protein segments found in different architectures

**DOI:** 10.1101/2020.12.22.424031

**Authors:** Rachel Kolodny, Sergey Nepomnyachiy, Dan S. Tawfik, Nir Ben-Tal

**Affiliations:** University of Haifa, Department of Computer Science, Haifa, Israel; Weizmann Institute of Science, Department of Biomolecular Sciences, Rehovot, Israel; Tel Aviv University, George S. Wise Faculty of Life Sciences, Department of Biochemistry and Molecular Biology, Tel Aviv, Israel

## Abstract

The vast majority of theoretically possible polypeptide chains do not fold, let alone confer function. Hence, protein evolution from preexisting building blocks has clear potential advantages over *ab initio* emergence from random sequences. In support of this view, sequence similarities between different proteins is generally indicative of common ancestry, and we collectively refer to such homologous sequences as ‘themes’. At the domain level, sequence homology is routinely detected. However, short themes which are segments, or fragments of intact domains, are particularly interesting because they may provide hints about the emergence of domains, as opposed to divergence of preexisting domains, or their mixing-and-matching to form multi-domain proteins. Here we identified 525 representative short themes, comprising 20-to-80 residues, that are unexpectedly shared between domains considered to have emerged independently. Among these ‘bridging themes’ are ones shared between the most ancient domains, e.g., Rossmann, P-loop NTPase, TIM-barrel, Flavodoxin, and Ferredoxin-like. We elaborate on several particularly interesting cases, where the bridging themes mediate ligand binding. Ligand binding may have contributed to the stability and the plasticity of these building blocks, and to their ability to invade preexisting domains or serve as starting points for completely new domains.

## Introduction

Over the course of 3.7 billion years of protein evolution protein segments of varying lengths mutated, duplicated, and recombined [1–5]. Contemporary proteins may hold hints to these ‘historical’ events. A likely scenario is that they evolved by duplication and fusion of short polypeptides with at least marginal stability, and weak biological functionality, sufficient for their preference over random alternatives. By mining protein databases [6–9], one can computationally search for traces of the evolutionary events that shaped the current protein universe, such as mutations, duplications, and recombinations of short protein segments (e.g., [4, 10–15]). Convergence is also a scenario that may result in sequence similarity. But because sampling a specific sequence (even as short as a few dozen residues) from the vast number of possible sequences is an extremely low probability event, when sequence segments of sufficient similarity are detected, common ancestry (homology) is the more likely scenario [16]. In practical terms, there are accurate, fast, and sensitive sequence aligners (e.g., HHSearch [17] or HMMER [18]) to identify similarities indicative of common descent. Most of the observed homology among current-day proteins may reflect relatively recent events. Nonetheless, there is also hope of finding protein segments that are relics of primordial, ancestral peptides that gave rise to what is now two or more separate lineages. These segments would typically comprise conserved and functionally critical parts of contemporary proteins [19, 20]. In the words of Eck and Dayhoff in their seminal 1966 paper, this is due to “natural selection which severely inhibits any change to an (ancient) well-adapted system on which several other essential components depend” [1].

The most widely recognized form of shared segments are protein units of approximately 100 residues called domains [9, 11, 21–23]. The definition of what is a domain varies [24], emphasizing structure [25] or sequence [26]. In the structure-based definition, domains are segments that fold independently to standalone (and even globular) entities [27]. Using this definition, protein space includes overall different domains with similar substructures [28, 29], which might reflect biophysical constraints on the protein chain [30–32]. In the alternative definition, emphasizing sequence, domains are commonly found protein segments that share significant sequence similarity with each other [26]. Indeed, using this definition reveals domains as evolutionary entities found in different combinations: or, equivalently, protein space has many instances of overall different protein chains with shared domains [11, 33]. The domain databases use the latter definition and group domains of the same evolutionary lineage [9, 26, 34–36]. When studying protein evolution, the initial focus was on standalone domains, because unlike short segments that cannot even fold, these can readily serve as evolutionary building blocks. This description, however, makes one wonder about the emergence of domains, and whether segments that are smaller than a domain, yet cannot fold or function on their own, played a role in that.

Indeed, sequence similarity among segments shorter than domains has also been described [4, 12, 13, 16, 37–40]. In fact, we have observed that the number of statistically significant similar segments increases with the decrease in their length (number of amino acids) [4]. Presumably, the proteins in the ancient protein universe were shorter, and the long period of time that passed offered their sequences many “copy-paste” opportunities. Consequently, short segments that show meaningful sequence homology are candidates for such ancient segments. Similarity between short segments can be classified into two types. The first includes series of repeated or amplified (similar) copies of a given segment in the same protein chain [5], suggesting emergence by duplication and fusion. Indeed, repeated segments can be identified from the internal symmetry in the sequence of the protein chain. The example that Eck & Dayhoff identified early on, was a short repeating segment in ferredoxin binding an iron-sulfur cluster [1] (see [41] for a retrospective view of this discovery). More recent examples include the ancient double β-hairpins and longer elements identified in the Outer Membrane Beta Barrels (OMBBs) [15, 42, 43], and the repeating β-blade forming β-propellers [44, 45]. The second type comprises homologous segments found in different contexts, namely in proteins that are deemed to have no common evolutionary origin. Prominent examples are the KH-motif [16, 46], the short segments with the same function in different SCOP folds [47], and the Fuzzle database [48]. Most relevant to this study is Alva, Söding, and Lupas’s curated set of 40 ancient segments that are shared among domains of different SCOP folds [49].

In a previous study we systematically documented similar protein segments that are shared between different proteins, referring to them as ‘themes’ and to the individual occurrences within each theme as ‘variations’ [4]. Here, we take advantage of these themes to describe a set of yet unknown ‘bridging themes’, namely homologous protein segments that are found in different sequential and structural contexts. The challenge in constructing such a set is that most similarity is detected among homologous domains, rendering the shared origin of such themes trivial (as these domains share common ancestry). To avoid these, we look only for cases where the variations of a given theme are found among domains whose overall sequences and structures are different, thus excluding shared ancestry of the entire domain. That a theme is shared between two current-day domains that are thought to have evolved independently suggests that this theme may have played a significant role in the emergence of these domains. Specifically, a shared theme may reflect common ancestry, although its precise role may vary. Assuming that modern proteins evolved from short polypeptides, a single ancestral fragment could extend by accretion or fusion to different other segments, and ultimately give rise to two different domains each of a different fold. Alternatively, a segment can be coopted from a preexisting domain, and fused to another segment, or duplicated to generate a new fold. At this stage, these two scenarios cannot be distinguished. We thus dub these themes, bridging themes, since regardless of the precise scenario, that they are currently found within the seed of two fold(s)/domain(s) that are deemed independent evolutionary lineages attests to their ability to fit in different environments.

To detect these bridging themes, we search for sequence similarity in a (non-redundant) set of domains that are classified as evolutionarily distinct [9]. In addition, we verify that beyond the shared theme, the rest of the domain sequences are not homologous. We find 525 such bridging themes, spanning 73 different folds, including the most ancient, pre-LUCA enzyme folds Rossmanns, P-loops, TIM-barrels, and Flavodoxins [50, 51]. The identified themes uncover many previously unknown potential evolutionary relationships, including ones that relate these ancient folds to each other. In approximately half the cases, the context change is also accompanied by significant alteration in the structure of the theme itself.

## Results

### Detecting bridging themes

We identify themes shared between non-homologous protein domains – cases where similar protein segments are found in two different contexts. More specifically, the segments should be similar to each other because they are detected using an adequately low HHSearch [17] E-value, and their similarity is verified by a high sequence alignment score. Yet their contexts are different as judged by structure- and sequence-based criteria. The structure context is deduced from the ECOD classification of the domains. The five-levels of the ECOD hierarchy, the so-called A.X.H.T.F groups, classify domains based on their structure and shared evolutionary origin [9, 52]. The top A (architecture) level does not indicate evolutionary relationships but rather, is based on the secondary structure content. The remaining levels of the hierarchy, from the X (possible homology) level downwards group domains based on presumed common ancestry. Domains with the same X classification (denoted X-groups) designate a distinct fold, and are possibly of common origin, yet more evidence is needed to establish whether they did in fact descend from a common ancestor. The next levels – H (homology), T (topology), and F (family), group domains with increasing levels of overall sequence and structural similarity, thus clearly indicating common ancestry [52]. To be conservative, we focus on themes shared between different X-groups, i.e., between domains that emerged independently even by the most lax ECOD definition of independent evolutionary lineages. The sequence context was also examined by applying, in addition to the above ECOD-classification-based filter, another filter based on sequence similarity. We look for cases where the best possible alignments of the domain segments flanking the shared theme (before the theme or/and after it) are poor, as these are indicative of a different sequence context.

Figure 1 illustrates the search process that we use to detect bridging themes. Using HHSearch, we search for shared themes in an ECOD database of domains that has been reduced in redundancy to 70% sequence identity (see Methods for details). Figure 1A shows this search for one instance, highlighting two domains in the database (domains 1 and 2) with segments that match the query with low E-values indicative of homology (< 10-^3^). We then select only those pairs of domains that belong to different ECOD X-groups. These pairs are broken to three parts (Figure 1B): the recurring part, the part before it, and the part after. We align the matching recurring parts to each other using a local (Smith Waterman (SW)) or a global (Needleman Wunch (NW)) aligner. Figure 1C summarizes the properties that we verify prior to including a pair in the set of curated bridging themes, that comprises 525 themes, spanning 73 different ECOD X-groups, or folds. The structure context is different, and the sequence context differs because the SW alignment scores of the parts before and after is low, and yet the shared theme is of common origin.

**Figure 1:**
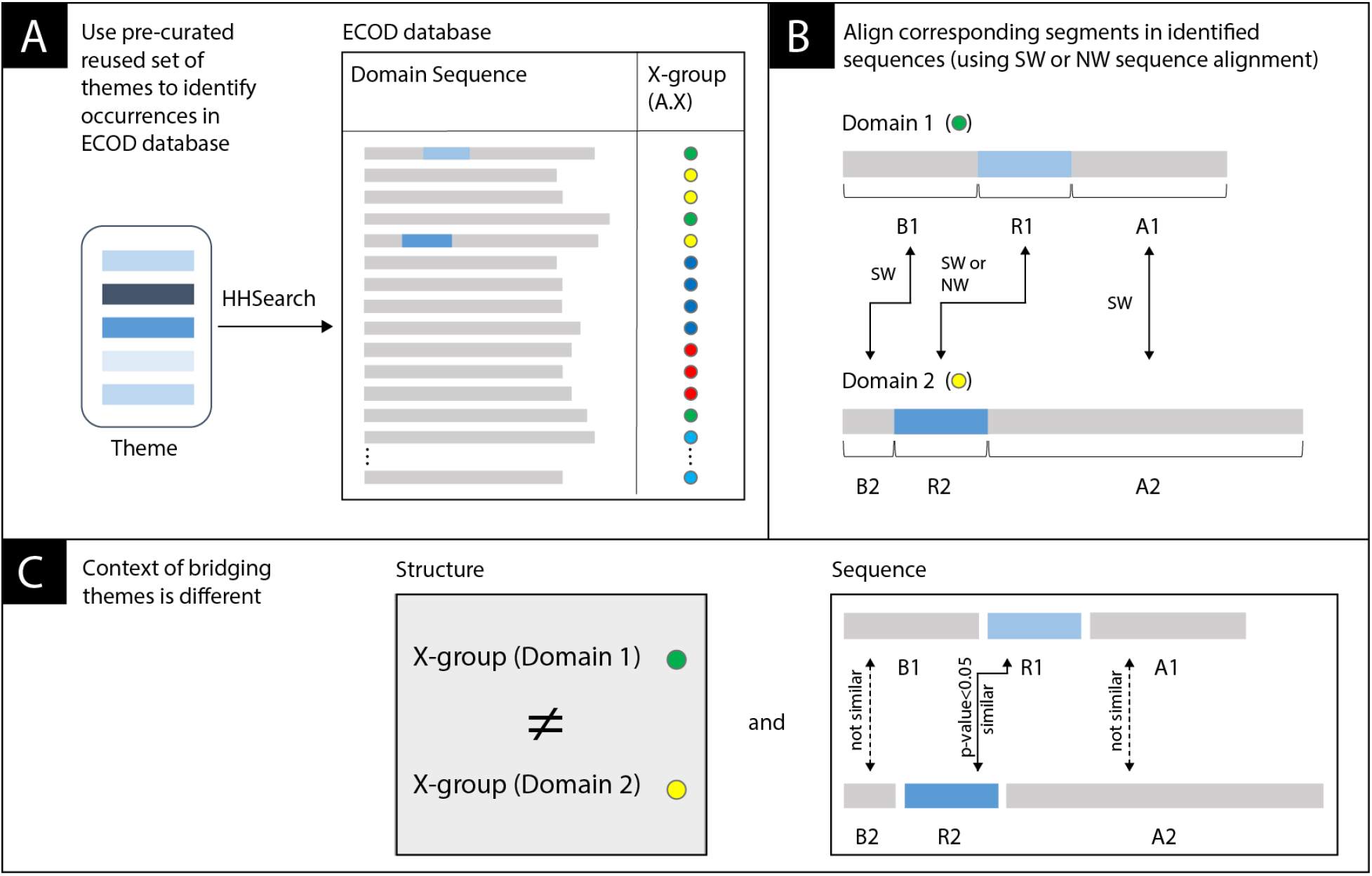
Overview of the process of identifying bridging themes. (A) We rely on a pre-curated set of 12,769 themes [4](one theme is represented here by shades of blue), and use HHSearch to search for segments in the ECOD domains database that exhibit sequence homology. Each ECOD domain is characterized by its X-group (a unique number), represented here by a colored dot. (B) Themes that appear in two domains each belonging to a different X-group (domains 1 and 2) are further analyzed by calculating the optimal sequence alignments of three segments: An alignment between the matched recurring parts (the themes, R1 vs. R2) using the local Smith-Waterman or the global Needleman-Wunsch algorithms, and local Smith-Waterman alignments between the segments before (B1 vs. B2) and the segments after (A1 vs. A2) the theme. (C) Overall, we search for events in which the context of the shared theme differs: the structure context differs if the domains are from different X-groups, and the sequence context differs if the local alignments (B1 and B2) and (A1 and A2) have low similarity.

### General properties of the bridging themes set

Figure 2 (A, B) shows the distributions of the length, and of sequence identity, or similarity, of the bridging themes in our set. For completeness, Figure 1S relates these measures and E-values, to one another. As dictated by our search procedure, the themes are longer than 20 residues (save 10 cases, where the alignment results in fewer residues), and because we search for similarities between domains of average length of 100 amino acids, the themes are generally shorter than 80 residues, with a mean length of ~49. The high mean sequence similarity of ~64% and mean sequence identity of ~30%, as well as the low P-values of the alignment scores, all indicate that these themes likely reflect common ancestry. In contrast, the segments before and after the bridging theme fail to show significant homology. Specifically, we align the corresponding parts before and after the recurring segments using a local SW aligner. In 85% (444) of the pairs, either the alignment is too short (less than 20 residues), or the best alignment results in sequence identity lower than 25%. For the segments after, in 88% (464) of the pairs, either the alignment is too short (less than 20 residues), or sequence identity is lower than 25%. In the remaining cases (which are left in the set but not described), the local SW sequence alignment in the part before or after is not significantly worse than that of the shared theme part. However, because this is a local alignment, it is not contiguous with the recurring part. It is possible, however, that the (evolutionary linked) recurring segments are longer (and detectable by allowing additional longer gaps).

**Figure 2:**
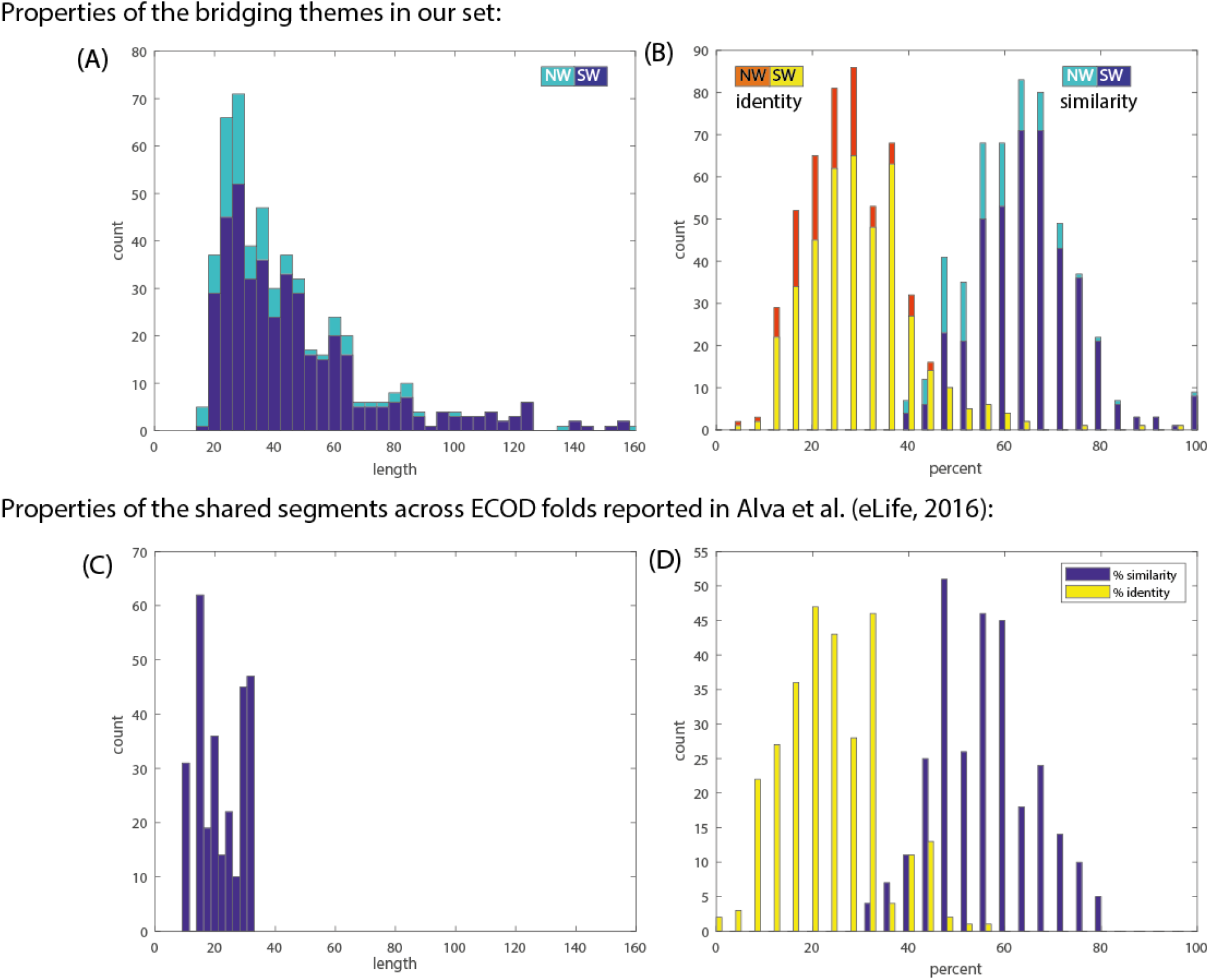
The cumulative properties of the bridging themes in our dataset vs. the fragment set of Alva et al. [49]. The distributions of length (A) and sequence identity/similarity (B) in our dataset. By design, our themes are longer than 20, and their mean length was found to be ~49 residues. The mean sequence similarity is ~64% and the mean sequence identity is ~30%. The distributions of length (C) and sequence identity/similarity (D) for the pairs of ancient fragments in the set of Alva et al. Shown is the subset of all pairs that span two ECOD X-groups (n = 286). The mean length of the recurring segment in this set is ~22 residues, the mean sequence similarity is ~56% and the mean sequence identify is ~24%. In general, the two datasets have similar criteria for selecting segments of shared origin, although our dataset is somewhat more conservative, and accordingly our themes are longer and with higher sequence similarity/identity.

The chains covering our themes include 121,749 residues in total, out of which 31,085 comprise the bridging themes. Figure 2S(A) compares the percent of different classes of secondary structure, and figure 2S(B) compares the percentages (i.e., a normalized histogram) of the solvent accessibility values. In both cases, the distributions are similar, i.e., the themes are not unique in their secondary structure nor their solvent accessibility.

### Structural similarity of the variations of the bridging themes

We did not restrict our search to cases where the variations of the bridging themes are structurally similar (i.e., within the two domains where the themes are found). Figure 3 shows the distribution of their RMSD (over the matching C-alpha atoms) and these indeed vary. The average RMSD over the whole dataset is 8 Å. Using a 6 Å RMSD threshold, the structures of the recurring themes in 39% (203) of the pairs are similar, and 61% (322) are not. Because RMSD is sensitive to outliers, and hence may be overestimating the variations, we used additional measures to quantify the structural similarity: We structurally aligned the matching domain segments with TM-align [53]: Figure 3SA,B show the histograms of TM-scores and RMSDs of the aligned residues. TM-scores are on a 0-1 scale, where scores > 0.5 indicate of the same fold and <0.3 corresponds to random structural similarity. In our bridging themes set, 30% have a score < 0.3, and only 25% have a score > 0.5. Figure 3SC-F shows the histogram of percent agreement of secondary structure assignment (by DSSP), dRMSD (distance RMSD), and percent contact map change of the aligned residues (see Methods for details). These measures also indicate that our bridging themes set includes pairs with different structures. The structural variation is consistent with the sequence homology being the outcome of common ancestry, rather than convergence due to structural constrains. It appears that one of the qualities of the bridging themes is that their structures can vary in accordance with their different contexts.

**Figure 3:**
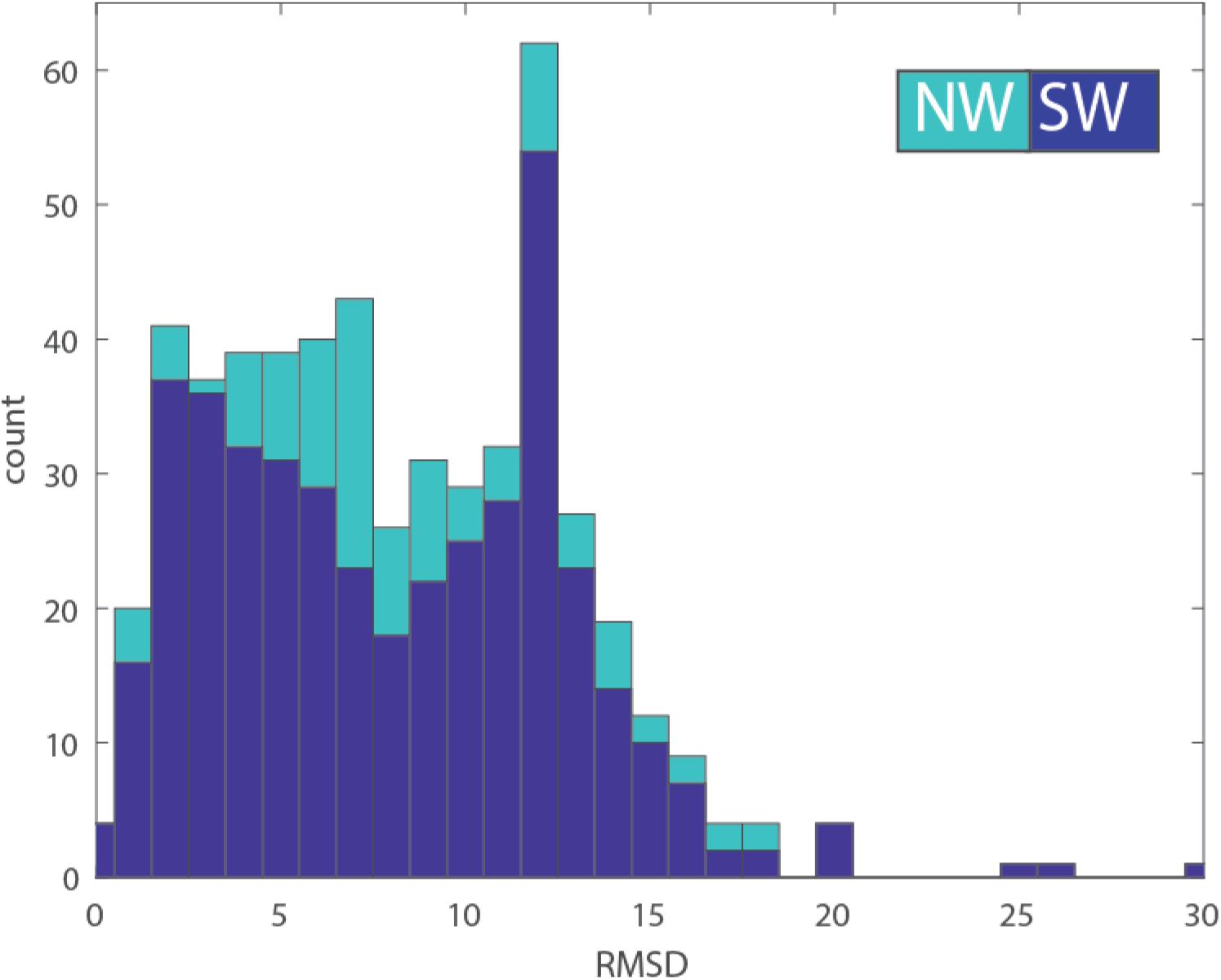
The histogram of RMSD values between the variations of the bridging themes in our dataset. The values for the Smith-Waterman (local) alignments are shown in blue, and for the Needlman-Wunch (global) alignments in light-blue. The global average is ~8 Å RMSD, yet the histogram appears to be a mixture of two distributions, one of structurally similar themes (characterized by lower RMSD values), and one of structurally dissimilar themes in spite of their high sequence similarity. Using a 6 Å RMSD threshold, 39% of these sequence variations are structurally similar, and 61% have different conformations. When fitting a mixture of two gaussian distributions we find that 28% of the pairs of variations that share the same theme also share a similar conformation (averaging ~3.3 Å) and 72% do not (averaging ~9.8 Å).

### Enrichment of binding residues within the bridging themes

For approximately half of the domains in our dataset, we can identify a binding function from their PDB structures. We identify binding residues (and the domains that include them) in one of two ways: (1) residues within 4.5 Å of a ligand, and (2) residues listed in the BioLip database [54] (see Methods for details). Because the bridging themes cover only some of the domain’s residues, binding residues may be either within the theme, or not. Had the binding residues been chosen uniformly and at random from all domain residues, we would expect that (on average) the proportion of binding residues in a bridging theme out of all binding residues is the same as the relative proportion of bridging theme residues in the domain. Figure 4S shows that when comparing these ratios, there are many cases above the diagonal line, i.e., the binding residues are more likely to be within a bridging theme than what is expected from the lengths of these themes. The total number of residues in the binding domains are 30822 (BioLiP dataset) and 38724 (4.5 Å dataset); the total number of residues in the themes in these domains are 11533 (BioLiP) and 14135 (4.5 Å). The ratio between the two is 0.37 for both sets. The total number of binding residues within these domains, is 2400 (BioLiP) and 3843 (4.5 Å); the total number of binding residues that are in a bridging theme is 1321 (BioLiP), and 2062 (4.5 Å). The ratio between the two is 0.54-0.55, which is larger than 0.37, suggesting that bridging themes are more likely to relate to function compared to their flanking segments. However, it does not preclude other structural roles of bridging themes, especially because ligand-binding involves many amino acids beyond those that directly mediate the interaction with the ligand.

### Network views of the bridging themes

We detected 525 instances of bridging themes among 73 different X-groups belonging to 17 (of the possible 20) different architectures (A-groups). Table 1S lists these X-groups, and the number of domains in each. The number of members in the X-groups varies between 1 and 69 (ECOD id 2007, flavodoxin-like), with an average of 7. The X-groups are identified by their ECOD id: e.g., 101 for HTH, or 2004 for the P-loops. This set includes all-alpha, all-beta, alpha/beta, alpha+beta, and mixed alpha/beta and alpha+beta, suggesting that bridging themes are shared throughout the entire protein universe. We organize the bridging themes as networks in two-levels. Figure 4 shows the first, overview level. Nodes represent ECOD X-groups, colored according to their ECOD A-group classification, and edges connect the two different X-groups among which the detected theme is shared. We find themes shared between X-groups from almost all class combinations, and in particular relating alpha/beta and alpha+beta proteins to either all-alpha or all-beta ones. More strikingly, some themes are shared between all-alpha and all-beta proteins. As X-groups contain many domains, a pair of connected X-groups, i.e., an edge in the overview network, may represent more than one instance; an instance being a shared theme relating one of the possible pairs of non-redundant domains (one from the first X-group, and the other from the second; see Methods for details). Thus, for each edge in the overview network, we organize all its instances as a separate (nested) network. Nodes in the nested network (e.g., Fig. 5) represent the domains from the two X-groups, and edges in the nested network connect domains with variations of the same theme. A nested network may include alternative representatives from the 70%NR dataset from different X-groups, that are similar to each other within each X-group (e.g., Fig. 5). We use Cytoscape [55] / CytoStruct [56] to visualize the networks; the Cytoscape session is in the supplementary material, and online^1^.

**Figure 4:**
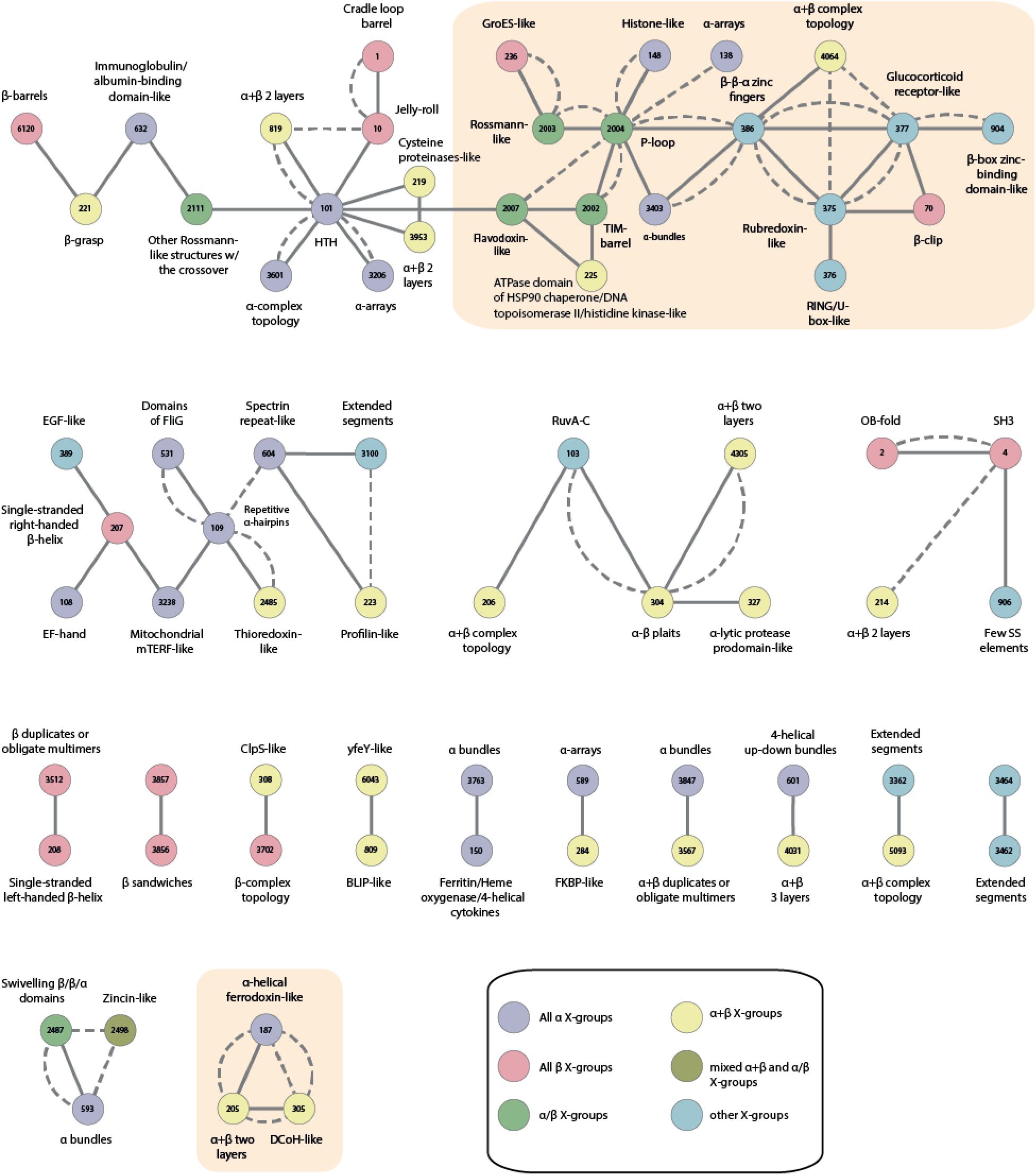
The overview network representing the 73 ECOD X-groups with bridging themes that are shared between them. The ECOD X-group number is listed in the node, and the corresponding name is listed next to it. The colors correspond to the X-group classes (see legend). Edges connect pairs of X-groups that share a common theme. The shared themes were found either by aligning the relevant segments in the two domains using a local (SW) or a global (NW) alignment. If the optimal local (SW) alignment of the shared theme that is longer than 20 residues, the edge is a solid line, otherwise, it is a dashed line. Connected components that are further discussed are highlighted with yellow background. The connected component on the upper row is further described in Figures 5 and 1S, and the connected component in the bottom row includes the examples in Figures 6 and 7.

**Figure 5:**
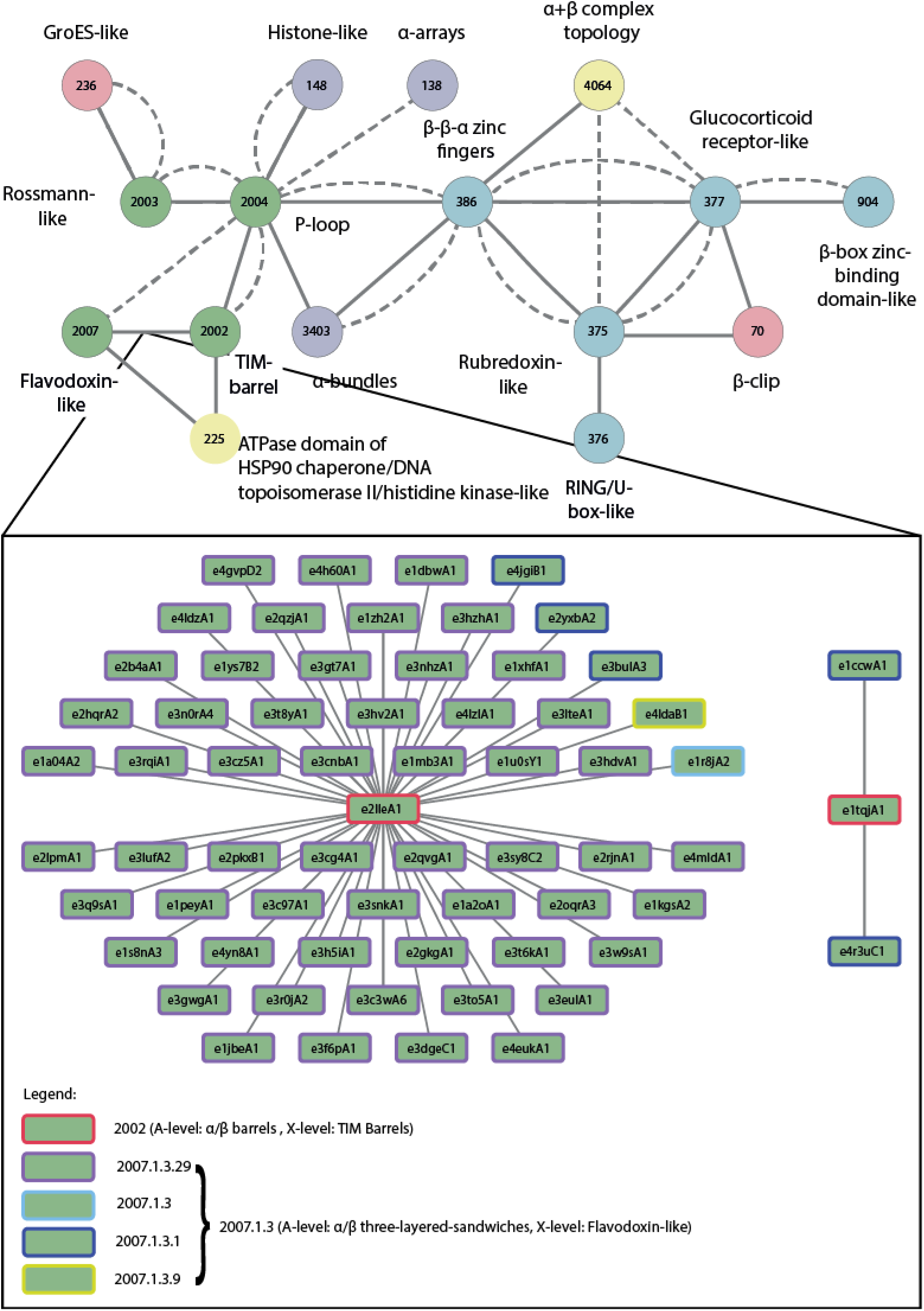
An example of the nested network describing the domains and their shared themes relating ECOD X-groups TIM-Barrels (2002) and Flavodoxin-like (2007). The nested network (lower panel) expands on an edge between these two X-groups in the overview network (a snippet of this network is shown in the upper panel). Within the nested network, each domain is described by a node, and edges connect pairs of domains, one from each X-group. The nodes’ background is colored by their A-level classification using the same color-coding as in Figure 4 (in this case, green, as they are all alpha/beta domains). The nodes’ boundaries are colored to differentiate their lower-level classifications, with arbitrarily chosen colors. Most of the shared themes are between the same TIM-barrel domain e2lleA1 and various Flavodoxin-like domains. However, there are also shared themes between another TIM-barrel domain, e1tqjA1, and two Flavodoxin-like domains.

The most extensively connected component in the overview network (Figure 4, uppermost) relates five alpha/beta X-groups (Rossmann-like, Rossmann-like w/crossover, Flavodoxin-like, a P-loop domains-like, and a TIM barrel), seven all-alpha X-groups (alpha bundles, two alpha arrays, Histone-like, alpha complex topology, immunoglobulins, and HTH), five all-beta X-group (GroES-like, jelly-rolls, cradle-loop barrels, and beta-clips), six groups of alpha+beta complex topology (alpha+beta 2 layers, beta-grasp, Cysteine proteinases-like, and ATPase domain of HSP90 chaperone/ DNA topoisomerase II/ histidine kinase-like), and five X-groups of few secondary structure elements. Interestingly, many of these, and the alpha/beta X-groups in particular, are considered to have been present in the LUCA [49, 57–59], suggesting that we traced events relating domains that evolved particularly early and that are not detectable by global domain sequence similarity. Figure 5S zooms in on part of this connected component to show examples of a shared theme for some of the pairs of connected X-groups. The most gregarious X-groups in this subset in terms of shared themes are the P-loops (id 2004) and the HTH (id 101). That they share themes with many other X-groups is consistent with previous estimates that they are ancient [59].

Figure 5 shows an example of the network of the themes shared between the Flavodoxin-like (2007) and the TIM-barrel (2002) X-groups; this is the network nested in the edge connecting the nodes 2007 and 2002. The domains in these two X-groups cluster into two connected components. The first is star-like, with the TIM-barrel domain e2lleA1 at its center, connected to no fewer than 60 different Flavodoxin-like domains. The single domain protein e2lleA1 (PDB ID 2lle) was engineered by Höcker and co-workers as a ‘copy-paste’ instance between these two X-groups [60]. That we detect this artificially designed protein validates our approach. In fact, because we allow sequence variability in the theme, we identify different Flavodoxin domains, similar (at least in part) to the one used in the design of 2lle. The domains belong to the ECOD T-group 2007.1.3 (Flavodoxin-like/Class I glutamine amidotransferase-like/CheY-like): 55 of them in the F-group 2007.1.3.29 (Response_reg), 3 in 2007.1.3.1 (B12-binding), one in 2007.1.3.9 (OKR_DC_1_N_like); e4ldaB1 is unmapped. Interestingly, the second connected component in Figure 4 reveals themes shared between the naturally occurring Flavodoxin domains e1ccwA1 and e4r3uC1 (both, F-group 2007.1.3.1 B12-binding), and the TIM-barrel domain e1tqjA1 (see Figure 6S for more details), suggesting an evolutionary event independent of the artificial design.

### Relation to Alva et al.’s ancient fragments

Alva et. al. recently curated a set of 40 ancient protein fragments [49]. Their set covers all previously documented cases as well as instances that were not known before. We downloaded their set and assigned the ECOD classification to their fragments. Their set includes 40 fragments, where each fragment is described as an MSA of sequence segments from different proteins. 23 of these fragments span different ECOD X-groups. The number of sequences in the MSAs of each of their fragments varies from 2 (e.g., fragment #30 in their set) to 20 (fragment #1 in their set). For the purpose of comparing to our set of bridging themes, we consider a subset of sequence segment pairs that are in the same fragment, i.e., are aligned to each other in the MSA of that fragment, yet the ECOD X-groups of the domains of these segments differ; there are 286 such pairs in their dataset. For every such pair, we measured the number of aligned residues, and percent similarity and identify, after aligning the reported segments with the global NW algorithm. Figure 2(B, D) shows the distribution of these values. Compared to our set of bridging themes, their ancient fragments are shorter, between 9-33 residues (mean of ~22) vs. 16-380 (mean of ~49) in ours. The average sequence identity (24%) and similarity (56%) in their set is also lower than in our set (30% and 64%, respectively).

Figure 7S shows an overview network derived from the abovementioned 23 fragments in Alva et al.’s set that are classified to different ECOD X-groups (44 X-groups in total). Only 17 X-groups, and 3 pairs of X-groups, are found in both sets – ours and Alva et al.’s – (Figures 7S and 8S). That the overlap between the results is relatively small and is due to the different search strategies used. Our strategy makes explicit use of the expectation that the segments are themes, i.e., commonly used parts, as elaborated in the “Comparison to Alva et al.’s paper” section in Methodology.

### Examples of newly identified bridging themes

For concreteness, we briefly describe only some of the bridging themes. A notable evolutionary link depicts a theme shared by ECOD domains e1nekB1 and e2pmzS1 of the alpha-helical ferredoxin-like (187) and DCoH-like (305) X-groups, respectively (Figure 6). The structures are positioned so that the shared segments are superimposed based on the sequence alignment, although as can be seen they vary in structure (for clarity, the individual structures are shown on the sides; Figure 6A). The sequences of these two variations of the same theme are homologous (47 aligned residues with 34% sequence identity and 75% similarity; Figure 6B), yet their sequence context, namely the sequence segments before and after the shared theme, are not (Figure 6C). Also, not only are the overall structures of the domains different, but even the structures of the theme itself are quite different (optimal C-alpha RMSD is 9.7 Å). Although detected in two evolutionary distinct ECOD X-groups, both variations bind an F3S iron-sulfur cluster, further corroborating their shared evolutionary origin. Moreover, upon optimal structural superimposition of the two variations, their ligands reside in the spatial vicinity of one another (Figures 6A and 6D). Interestingly, even though the two variations share the same F3S ligand, their binding modes are somewhat different: F3S binding is coordinated by four cysteine residues in e2pmzS1 versus three cysteine residues and a serine in e1nekB1 (Figure 6C). Despite these differences, and the different structure of the shared theme in the two domains, two of the cysteine residues are well aligned. Accordingly, ConSurf [61] evolutionary analysis shows that the four residues that mediate F3S binding in both e1nekB1 and in e2pmzS2 are highly conserved among the homologues of the two respective proteins (1nek and 2pmz; Figure 6C). It is noteworthy that the two variations of the theme differ from each other in their conservation pattern. The e1nekB1 variation is much more evolutionarily conserved (among 1nek’s homologues) than that of e2pmzS2, reflecting the effect of years of evolution in different context: succinate dehydrogenase, where the iron-sulfur cluster is essential for enzymatic function vs. RNA polymerase, where it mostly plays a supporting role in stability and folding [62].

**Figure 6:**
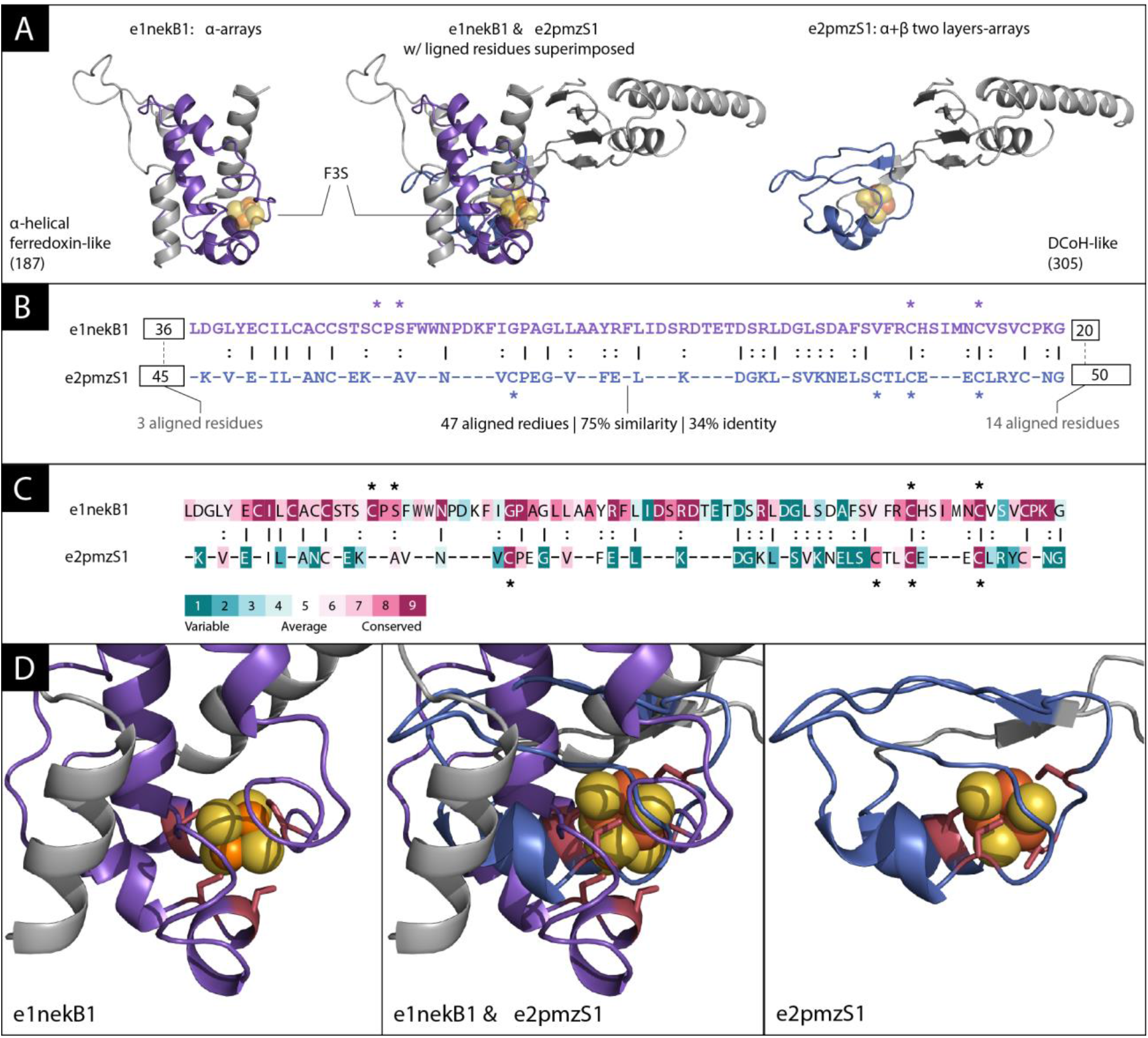
An F3S-binding theme shared between domain e1nekB1 from X-group 187 and domain e2pmzS1 from X-group 305. (A) The overall structures of these two domains is different: e1nekBa (left) is an alpha-helical ferredoxin-like from the all-alpha class, and e2pmzS1 (right) is a DCoH-like from the alpha+beta class. In the middle, are the two structures in which this theme is found, in the best possible superimposition of the 47 aligned theme residues, and their two corresponding F3S clusters shown as orange (iron) and yellow (sulfur) spheres. We see that the iron-sulfur clusters reside in equivalent locations. (B) When aligning the shared theme (magenta) there are 47 equivalent residues, with overall sequence similarity of 75%, and sequence identity of 34%. The residues binding the iron-sulfur (F3S) clusters are marked by asterisks: in e1nekB1, there are three cysteine residues (C159, C206, C212) plus a serine (S161), and in e2pmzS1 there are four cysteine residues (C183, C203, C206, C209). The sequence context of the two variations of this theme are different: The best local alignment of the ~40 residues before the theme has only 3 aligned residues, and the sequences after the theme share 14 aligned residues (of 20 residues in e1nekB1 and 50 in e2pmzS1). (C) ConSurf evolutionary analysis of the corresponding proteins (1nek and 2pmz) and their 150 respective homologues. As can be seen, the binding residues are highly conserved (3 with maximal conservation score 9, and one with conservation score 8 in both proteins). (D) Zooming-in on the domains’ iron-sulfur clusters. The two structures superimposed in the middle, and each one separately on both sides. The cluster binding residues are shown in red and their side-chains are shown as sticks.

Figure 7 shows an example of a theme shared between domains from ECOD X-groups 187 an alpha-helical ferredoxin-like (α-arrays ECOD A-group), and 205 a 4fe-4s ferredoxin (α+β-two layers ECOD A-group). Between these two X-groups, we found fifteen representative instances, connecting five domains in the 187 group, and thirteen domains in the 205 group. The figure shows one of these instances, where a theme of 32 amino acids appears in the α-helical e1kf6B1 and in the α+β domain e3mm5B5. Despite the high sequence similarity between the two variations of this theme, their structures are very different. Specifically, the secondary structure of the residues in the two contexts differ – there are only helices in the former and helixes surrounding a beta hairpin in the latter (Figures 7B, 7D). These differences are in accordance with the same theme being embedded in two very different sequence contexts (Figure 7B). Interestingly, both domains also bind a ligand – F3S in e1kf6B1 and SF4 in e3mm5B5 – encapsulated by some of the residues of the theme and positioned similarly with respect to it. Figure 7D zooms in on the region in space where the variations encapsulate the ligands, showing how well the ligands align to each other upon superimposition of their respective variations. The sequence of two variations of the shared theme align well with 63% sequence similarity and 38% identity (Figure 7B). However, the segment inserted in the middle of the theme differs between the two variations (Figure 7B): in e1kf6B1 there are 27 residues and in e3mm5B5 only three. Their difference indicates that both insertions may have taken place after the emergence of the theme. In e1kf6B1 the inserted 27 residues form a helical-hairpin, with many of its residues (180-192) exposed. Oddly enough, this insertion is in the folding core of the domain, suggesting that is has been key to the formation of the domain. Accordingly, homologues of e1kf6B1 (in the F-group 187.1.1.5) share this inserted segment, suggesting that it was one of the defining events in the diversion of this group of domains. The flanking segments do not align well at all: Before the matched theme there are 50 residues in e1kf6B1 and 22 in e3mm5B5, and yet the optimal local alignment matches only 8 residues. After the matched theme there are 28 and 9 residues, respectively, but the optimal local alignment matches only 2 residues. ConSurf’s evolutionary analysis of the two corresponding proteins (1kf6 and 3mm5) within the context of their respective homologues shows that the residues that mediate iron-sulfur cluster binding, and several other residues, are highly conserved (Figure 7C). Of these, the three cysteine residues in e1kf6B1 align with their equivalents in e3mm5B5, reflecting that despite different ligands (F3S vs. SF4), these two variations resemble each other in their binding modes.

**Figure 7:**
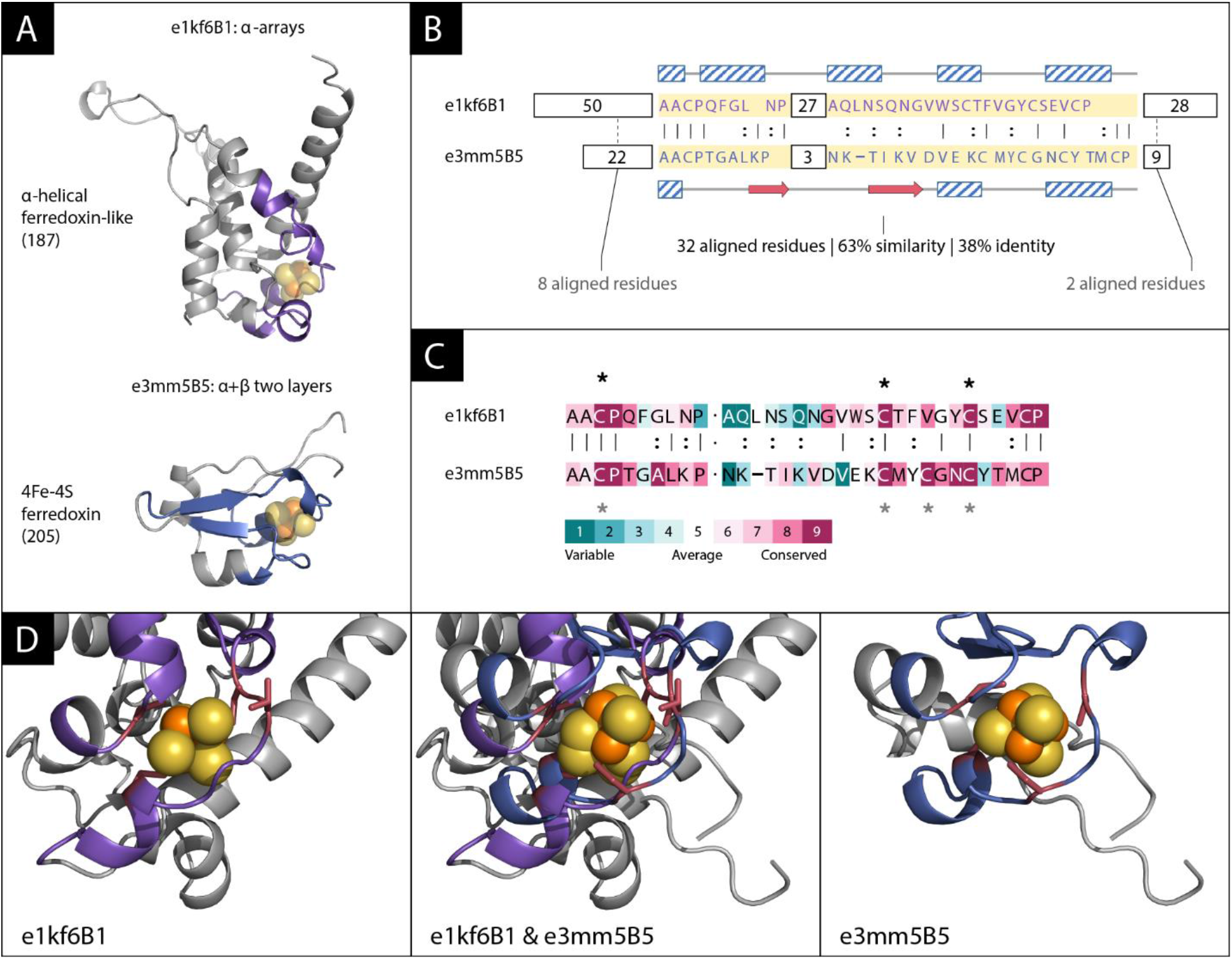
An iron-sulfur cluster-binding theme shared between e1kf6B1 (a ferredoxin-like domain of the all-alpha class) and e3mm5B5 (a 4Fe-4S ferredoxin domain of the alpha+beta class). Fifteen representative instances connecting five domains in the 187 group (e1e7pB1, e1kf6B1, e1nekB1, e3cf4A4, and e3vr8B1), and thirteen domains in the 205 group (e1fxdA2, e1hfeL2, e2fgoA1, e2gmhA1, e2v2kA1, e2wscC1, e2xsjB8, e2xsjD9, e2zvsA1, e3eunA1, e3j16B5, e3mm5B5, e4id8A1), are found between these two X-groups. (A) The structures of the two domains, including the recurring theme (magenta), are different. Two bound ligands (FS3 within e1kf6B1, and SF4 within e3mm5B5) are shown in atoms-spheres representation (iron in orange and sulfur in yellow). (B) While alignment of the shared theme suggest common ancestry (38% identity and 63% similarity over 32 residues), the secondary structures of the two variations differ: the α-helices are marked by diagonally patterned blocks and the β-strands by red arrows. Accordingly, the parts before and after the shared theme have different lengths and secondary structures, and cannot be successfully aligned. (C) ConSurf analysis of the corresponding proteins (1kf6 and 3mm5) and (150 of) their respective homologues show that the binding residues (marked with asterisks) are highly conserved. In e1kf6B1, C158, C204 and C210 were assigned a conservation grade of 9, and T205 (the resides in proximity to the cluster but may not be ligating it directly) a grade of 7. In e3mm5B5, all four cysteine residues (C220, C241, C244, C247) were assigned a grade of 9. (D) Zooming-in on the binding sites. The Iron-sulfur cluster is shown as orange-yellow spheres, the themes are colored in magenta, and the cluster binding residues in red the side-chains of the binding residues are shown in sticks. In the middle, the two structures are shown after superimposing the aligned theme residues, and on the left/right the structures are shown individually. Superimposing the aligned variations of the theme in the two domains (middle), results also in their F3S and SF4 ligands being well aligned.

Shorter versions of iron-sulfur cluster binding motifs were reported based on sequence and structure similarity searches [16, 63, 64]. In particular, the bridging theme of Figure 7 is closely related to fragment #18 in Alva et al.’s set, which connects X-groups 187 and 205. However, fragment #18 is about half the length. Two of the 14-residue segments that it includes are from the domain e2bs2B1, a homologue of e1kf6B1. The first segment overlaps with the part before the gap in our bridging theme (its last 5 residues align to the first 5 residues of our bridging theme), and the second segment overlaps with the last 13 residues of our bridging theme. One might argue that the high (more than 30%) sequence identities in the shared themes described in Figures 6 and 7 have emerged by convergence [63]. Indeed, the need to coordinate the cluster in a way that will enable proper electron transfer function imposes not only cysteines as the ligating (coordinating) residues, but also that these cysteines locate at certain distances from one another and with the right stereochemistry. We note, however, that ferredoxin-like domains that contain iron-sulfur clusters are highly abundant and span 16 ECOD X-groups (and likely more X-groups not annotated as such). Despite that, shared themes are detected only between few of these X-groups. That not all of the iron-sulfur binding proteins share the same theme suggest a common evolutionary origin rather than convergence.

Figure 9S shows a theme shared by ECOD domains e3ephA1 and e4dgwA1 of the P-loop (2004) and alpha-bundle (3402) X-groups, respectively. In this case, the structures of the shared theme of 26 residues (with 50%/28% sequence similarity/identity) are very similar in both domains (C-alpha RMSD = 1.38 Å) and they wrap, or enclose, similarly positioned Zinc ions. Zinc binding in 4dgw (an SF3a core protein) is coordinated by four invariant residues: C282, C285, H298, and H304 [65]. The four equivalent positions in the variation in e3ephA1 feature the exact same amino acids. In this case too, ConSurf analysis indicates that both sequences are evolutionarily conserved, each within the context of its domain, and so are the two cysteines and two histidines (assigned with the maximal ConSurf conservation grade). Structurally, this theme appears detached from the rest of the domain in both contexts, suggesting that it could be autonomously folded, perhaps stabilized by the bound ion. Thus, this theme may represent a case of cooption, namely of a segment taken from one protein being fused to another protein.

Finally, another tantalizing evolutionary link we have identified connects the two most ancient and diverse lineages of Rossmann-like and P-loop NTPases, and is analyzed in detail separately [20]. Similar to the above presented cases where the shared theme includes a key ligand binding motif, i.e., iron-sulfur clusters, this shared theme includes two key functional elements – a phosphate binding loop and an aspartate that binds the ligand’s ribose moiety in Rossmanns and the catalytic metal ion in P-loop NTPases.

## Discussion

Our systematic search for bridging themes, short homologous sequences shared between proteins that are assumed to have evolved independently, yielded 525 representative domain pairs, spanning 73 different folds, or ECOD X-groups. These themes have at least 20 residues, and the average percent sequence identity/similarity between their corresponding variations is high (30%/64% respectively), strongly indicating that these variations descended from a common ancestor. Organizing the representative examples as an overview network manifests the regions of the protein universe that they traverse. We find that the ‘alpha+beta’ (22 instances) are the most common, followed in descending order by ‘all alpha’ X-groups (20 instances), ‘all-beta’ (13), ‘others’ (11), and finally the ‘alpha/beta’ (6) and ‘mixed alpha+beta and alpha/beta’ X-groups (1). There are connections among X-groups of the same architecture (or A-class) [66], but also many that cross class boundaries and involving all class combinations.

The largest connected component in our overview network includes (among others) themes shared between five alpha/beta X-groups of fundamental importance: Flavodoxin-like, TIM-barrel, P-loop NTPase, Rossman-like, and Rossmann-like w/crossover. These groups share few notable features. They comprise the most ancient enzyme classes, and their founding function was phospho-ligand binding, and specifically binding of phosphorylated ribonucleotide ligands. Further, they bind the phosphate moiety at the N-terminus of a helix, and, with the exception of TIM-barrels, this helix always comprises the first helix (the binding element is usually described as a P-loop, and resides between the first beta-strand and first helix) [67]. Evolutionary linkages between some of these groups have been suggested. However, so far, only the link between TIM-barrels and Flavodoxins has been established [68]. Here, tangible links between all these folds are unraveled. The link between P-loop NTPase and Rossmann is of particular interest, and is discussed in detail in an accompanying manuscript [20].

The alpha/beta X-groups are interesting because they include many superfamilies [69], with various functions [70–72], and because they include domains that are considered ancient [49–51, 57–59, 73, 74]. There are other domain groups that are considered ancient in our overview network. For example, the cradle-loop barrel and the HTH [50, 51]. It was difficult to predict beforehand if the search for similar amino acid segments in protein space may reveal themes shared among ancient proteins because the signal for common-origin, namely sequence homology, is expected to diminish over time. However, this could be balanced by: (i) longer evolutionary time, which allows for more opportunities for the emergence of new folds by cooption of bridging themes, and (ii) by evolutionary pressure to preserve those segments that mediate key functions, rendering these also easier to detect. That we detected multiple shared themes among groups of likely ancient domains indicates that there may be cases in which these two effects dominate.

Our study complements the seminal study of Alva et al. who looked for ancient fragments in current day proteins, and identified a set of 40 short fragments that exist in different structural contexts [49]. We address a closely related aim, relying on the ECOD classification (rather than SCOP used by them), and focusing on identifying pairs of seemingly unrelated domains with a shared theme. To compare the results of these two efforts, we analyzed Alva et al.’s fragments in a framework like ours. Comparing the distributions of the lengths of the shared segments and percent sequence similarity/identity we see that the criteria for concluding that two protein segments have a common evolutionary origin are similar, but we tend to be more conservative with higher percent sequence identify/similarity and longer shared protein segments. Note, however, that in our dataset, we did not require that the sequence-similar variations also have similar structures, and indeed the structures are often different. Both studies rely on the state-of-the-art HHSearch sequence search engine [17], but the search strategies differ. Our study employs our previously curated dataset of themes [4] as “baits”, which we use to search for cases where variations of these themes appear in two or more different sequence and structural contexts. Using bait themes allows our search to “fish” sets of domains that are promising candidates, and even more specifically, to identify within the domains in these sets the evolutionary meaningful recurring segments, i.e., the shared themes. That a variation of the shared segments must be first detected as a theme of at least 30 residues (and with our thresholds), appears as a limitation of the method. Indeed, many of the cases that Alva et al. report and that we miss, are due to this. However, we believe that this added requirement focuses the alignment procedure on identifying evolutionary relevant cases. We expect that an ancient or bridging segment will be reused in protein space, and thus we also expect that it be detected as a theme.

The instances that we and Alva et al. [49] detect differ markedly. Only 17 ECOD X-groups and 3 specific pairs of X-groups that share the same theme, appear in both sets (out of the combined 100 ECOD X-groups). The cases in the Alva et al. dataset that we miss, are generally due to our additional requirements, e.g., minimal length, or that there is a matching bait theme. Nonetheless, in different regions of the protein universe, we find recurring themes of similar characteristics. In other words, even using our conservative thresholds, our approach significantly expands the set of documented events of protein segments shared between domains that are considered evolutionarily unrelated. Of particular interest are additional themes that include the ancient and diverse alpha/beta X-groups. The high sequence similarity among the variations of these shared themes suggests that they have emerged from a common ancestor even though they are found in two contemporary domains that do not share an evolutionary origin in their entirety.

There are alternative explanations to protein segments in distinct domains that are similar to one another. Either they formed independently, and their similarity is due to pure chance or convergent evolution, or, as we try to argue here, they share common ancestry. It is hard to discern which is correct. If we can estimate the probability of forming these segments, and it is very low, we have a probabilistic argument that undermines convergent evolution. The lower the probabilities, the stronger the argument, e.g., when the segments are long, or enriched with rare amino acids. Here, we use E-values estimates of the state-of-the-art method HHSearch [17] to identify segments for which we have probabilistic support that they are related to the same bait theme. Also, we kept only alignments that are relatively long and with significant p-values, as these have additional probabilistic support. However, the probabilistic model is based on sequence and ignores possible dependencies that may be due to structure. Biophysical constraints, e.g., due to the polypeptide backbone and protein structure, significantly limit structure space [30–32], presumably also constraining sequence space. Independently formed proteins may converge to similar structures [75] and thereby also to similar sequences [76], leading to false-positive hits. Being aware of these issues, we use the E-values and p-values to identify interesting cases, rather than as statistical estimates. In this context, it is noteworthy that because this study involves many sequence comparisons one might wonder whether correction for multiple hypotheses testing is required. This would have been the case had we relied on the E- and p-values for statistical estimates. However, because we use them only as filter thresholds, the issue is moot. In summary, while it is safe to assume that most of the similarities detected are real, we cannot commit to each individual case.

For cases of shared ancestry, we now speculate about the turn of events that could have led to such current-day patterns. One possibility is that in the ancient past, a short ancestral theme existed on its own, i.e., without the segments that flank it in the intact, contemporary domain, perhaps bound to an ion or mineral (e.g., Figures, 6, 7, 2S, and [1]), a nucleotide [77], or RNA [5, 16]. Over time, it may have duplicated, the two copies diverged (albeit not beyond the level of detection), and protein segments accumulated before and/or after both variations. Because sequence expansion happened post-duplication, the two duplicates expanded independently of each other, leading to two different sequence contexts and often to two different folds. Furthermore, because the ancestral short theme is only a small part of the two otherwise different domains, the resulting overall structures are different, appearing today as two (or more) unrelated ECOD X-groups. The different context may in itself result in the very same sequence (theme) adopting a different structure [78–81], and indeed, the structures of theme variations often vary, including sometimes even different secondary structures (e.g., Figure 3; Figure 7).

An alternative scenario is that the theme existed within the context of a functioning protein domain. Then, this part of the domain (i.e., that theme) was coopted, duplicated and inserted into another protein domain (akin to copy-pasting), or alternatively, duplicated and fused to generate a repeat protein. That a protein segment can be grafted into another protein is supported by the protein design experiments which carried out such scenarios, including cases when the source and destination domains are of a different ECOD X-group classification [60, 68]. For themes that arose via the first scenario, one can deduce that they are ancient, and that their context (i.e., their flanking sequence segments) evolved to accommodate and extend these enclosed themes. In contrast, in themes that arose via the second scenario, selection would act to readjust the coopted theme to the new context. We currently cannot determine which (if any) of the identified themes followed the first scenario, and which (if any) was subject to the second one. Moreover, these scenarios are not mutually exclusive. That is, an ancient theme that acquired additional protein segments over time, may have been subsequently grafted onto yet another protein. Phylogenetic reconstruction may shed light on this question. Regardless of the evolutionary scenario, that the themes are shared between domains from different X-groups attests for their evolutionary plasticity.

The feasibility of these scenarios can be examined experimentally. To this end, the themes that appear to mediate specific functions would be most convenient. Showing experimentally that an isolated theme retains its function (e.g., metal, iron-sulfur, or cofactor binding), even if at low level (e.g. weak affinity), would support the feasibility of the first scenario. On the other hand, grafting a theme from one context, to a domain in another context would attest to the feasibility of the second scenario. A copy-paste event is a plausible explanation to the observed themes shared between the flavodoxin domains e1ccwA1 and e4r3uC1 and the TIM-barrel domain e1tqjA1, as supported by the experiments of Höcker and co-workers [60, 68, 82] (even though the domains e1ccwA1, e4r3uC1, and e1tqjA1 were not studied directly in this experiment). To take a step further, a variation of the zinc-binding theme described in Figure 2S could be experimentally inserted into a designed protein to introduce zinc-dependent regulation, or similarly, the iron-sulfur cluster binding themes outlined in Figures 6 or 7 could be added to a protein to endow binding of an iron-sulfur cluster ligand. In other words, the themes, and specifically, their reconstructed ancestral sequences, may be good candidates for protein engineering (in contrast with the contemporary variations, that may have lost the contextual agility [45, 83]). Themes with ligand binding function are particularly attractive candidates [77, 84].

## Prospect

Domains are considered central to protein evolution [27, 85, 86]. In single domain proteins, a primordial ancestor with promiscuous enzymatic activity could be the progenitor of a diverse family of proteins with various activities towards a multitude of substrates. For example, all contemporary TIM barrel and Rossmann domains can be traced back to their respective common ancestors [3, 19, 57]. Further, a contemporary multi-domain protein with a novel function may have evolved by concatenation of primordial domains with their respective functions [3, 11, 21, 27]. However, it is yet to be known how the domains themselves evolved from smaller protein segments [5]. Bridging themes may provide hints to this end, and some of them may report the building blocks from which today’s intact domains evolved. To fulfill this role, they would probably have to be at least marginally stable, and should provide some advantageous biochemical function to be evolutionarily selected over other peptides. The iron-sulfur cluster binding themes of Figure 6 and 7 are good candidates in both respects. Iron-sulfur clusters are ancient relatively stable minerals [87, 88], that provide diverse stereochemistry and catalytic opportunities [1]. Thus, an iron-sulfur cluster might have been a nucleus that bound amino acids and/or di- or tri-peptides that eventually elongated towards the emergence of an iron-sulfur cluster binding theme, as hypothesized for ferredoxin [1, 89]. In support of this hypothesis, the iron-sulfur cluster of the D dimerization subunit of the RNA polymerase (2pmz, Fig. 5) contributes to stability, and mutations of the four cysteine residues that mediate its binding lead to aggregation of the subunit [90]. By analogy to domains, variations of a theme could have presumably emerged from a common ancestor, which then gave rise to different functions. It has been recently proposed, for example, that a ‘theme’ comprising a beta-alpha-beta element may have given rise to both the Rossmann lineage (by virtue of binding FAD or NAD) and the ferredoxin one (by binding an iron-sulfur cluster [88]). That variations of the theme of Figure 6 bind both F3S and SF4 iron-sulfur clusters supports this hypothetical scenario. To push the domains analogy even further, just as multi-domain proteins may emerge by mix-and match of existing domains [11], domains might emerge by mix and match of two or more preexisting themes. Protein engineering experiments could be conducted to examine function alterations within a given theme as well as concatenation of themes to integrate several functions and yield intact domains.

## Materials and methods

The search carried out in this study is challenging because we look for domain pairs that satisfy two somewhat opposing criteria: (1) share short matching segments of similar sequences, and yet are (2) dissimilar in their overall sequence and structure. To tackle this challenge, we search the ECOD database using a set of previously curated themes [4] as “baits”, and look for themes that appear in two or more dissimilar domains. Themes are sub-domain recurring protein segments that we have previously identified from all-vs-all sequence alignments of (non-redundant set of) PDB proteins [4]. More specifically, we used HHblits [91] to calculate an HMM model for each recurring segment (adding sequences from uniclust30). The themes in the pre-curated dataset are these HMM models of the reused segments. The lengths of the themes in this curated set vary but are at least 30 residues. Relying on the pre-processing step of curating the set of themes, and using these as baits holds two advantages: (i) it restricts the search, and rather than considering all domain pairs it focuses on the pairs that have a segment similar to the same theme, and (ii) we can derive the matching short segments within these domains from the parts aligned to the bait theme. The advantages of this search method come hand-in-hand with its inherent limitation: shared parts must first be detected as a theme (of at least 30 residues). Our bridging themes were identified using a total of 261 bait themes. While most (136) baits identified exactly two domains (one in each X-group), some identified more than two domains. Figure 10S shows the histogram of the number of domains identified by our baits.

We used HHSearch (version 3.0.0) to compare a set of 12,681 previously curated themes to a 70% NR set of ECOD domains (43,830 domains in version develop210 and 43,281 in the updated develop263). We used the E-value threshold of 10-^3^ and coverage=0.85 to identify significant hits and find 22,381 ECOD domains that are similar to any of the themes (spanning 3,137 F-groups, 746 H-groups, and 646 X-groups). 1,698 of our themes are similar to ECOD domains that are in different X-groups. Because there is extensive redundancy among the themes, we next identify representative meaningful examples. For each such theme, and for each pair of X-groups X_1_ and X_2_ that it matches, we consider all *n_1_* domains it matches in X1 and all *n_2_* domains that it matches in X_2_; we consider the *n_1_×n_2_* potential domain pairs. For each domain pair, we calculate the SW alignments of the parts before and after, and the SW or NW alignment of the recurring part that matches the theme. When the local SW aligner finds an alignment of the recurring parts that matches more than 20 residues, we opt for this alignment. Because the local SW aligner may discard the beginning and the end of a segment if they do not improve the overall alignment score, it may result in alignments that match fewer yet more similar residues, and specifically, fewer than 20 residues. In the 104 such cases (~ 20%), we consider the NW alignment. If *n_1_×n_2_* equals 1, then we return this example. Otherwise, we normalize the scores of the matching recurring parts and the parts before/after (by subtracting the mean score and dividing by the standard deviation) and identify the two examples with the maximal difference between the (high) alignment score for the recurring part and the (low) alignment score for the parts before/after. For every pair of domains, if they share more than one recurring theme (regardless of the similarity among the themes that identified these domains), only the longest theme was kept.

Finally, for each aligned pair of protein segments, we calculate the properties of the alignment: the number of aligned residues, the percent identity, the percent similarity (using Blosum62), structural similarity measures, and the p-value. The p-value measures the significance of the alignment score with respect to scores of alignments of random segments (drawn from the same distribution). We estimate the parameters of their EVD (Extreme Value Distribution) from the scores of the alignments between the first segment and 1000 randomly chosen segments drawn from a multinomial distribution estimated from the second segment. We calculated the following structural similarity measures: (1) RMSD of the C-alpha atoms of the aligned residues after optimal superpositioning, (2) measures of structural similarity after structurally aligning the matching domain segments with TM-align [53]: TM-score and RMSD of the structurally aligned residues, (3) dRMSD (distance RMSD) of aligned residues, (4) percent agreement of secondary structure assignment. We derived the secondary structure assignments from pre-calculated DSSP files [92, 93] (chains 1we3, 3iyg, 3k1q, 4b4t, 4di7 are unavailable). (5) Percent change in contact maps of aligned residues. We follow the CASP convention [94] by which two residues are in contact if the Euclidian distance between their C-beta atoms (C-alpha in the case of Glycine) is below a threshold; we used a 9 Å and 11 Å as thresholds. The percent similarity is the percent of differing entries in the 0/1 *lxl* matrices representing the contact maps (where *l* is the number of aligned residues), i.e., the number of differing entries/*l*^2^.

To identify the domains with binding residues within 4.5 Å of a ligand we collected 129 relevant ligand codes. To identify these, we started with all ligand codes in the resNames field of the HETATMs and removed crystallographic additives (listed in [95]), ligands which do not appear in the BioLiP frequency file, modified residues (e.g., MSE, CME), HOH, UNL, and UNX. 288 of the domains include one of these ligands, and we identified all residues with an atom within the 4.5 Å distance from it as binding it. BioLiP lists binding residues in most (217) of these domains. The number of binding residues found by the two methods is very similar (0.86 correlation).

The data is organized as two Cytoscape [55]/CytoStruct [56] sessions: one for the overview network, and one with all the nested networks. The colors of the nodes are based on their ECOD A-group classification, grouped into structural classes [66]: all-α in blue (α-arrays, α-bundles, α-complex topology, α-superhelices), all-β in red (β-barrels, β-complex topology, β duplicates or obligate multimers, β-sandwiches), α+β in yellow (α+β complex topology, α+β duplicate or obligate multimers, α+β three-layers, α+β two layers), α/β in green (α/β barrels, α/β three-layered sandwiches), mixed α+β and α/β in yellowgreen, and others in cyan (extended segments, few secondary structure elements). In the downloadable session, a right-click on a nested-network edge opens (1) PyMOL [96] to show the structures of the two domains, with the themes highlighted and superimposed on each other, or (2) BioEdit [97] to show the aligned sequences. In the online version, a click on the edge opens the nested network, in which one can click on edge to download the PyMOL script, or shift+rleft-click to see the superimposed structures and the aligned sequences in the web-browser.

### Comparison to the set of [49]

We downloaded the set from the supplementary to Figure 3 in [49]. For each of the segments in the MSAs of their set of ancient fragments, we identified the ECOD domain of that segment and recorded its ECOD classification. In two cases, we shortened the segment by one residue because it fell on a domain boundary (1NT0 146-163, 1JX4 177-198). We focus on pairs of sequence segments that are in the same fragment, i.e., are aligned to each other in the MSA of that fragment, and that the ECOD X-groups of the domains of these segments differ; there are 286 such cases. We compared the properties of the two sets and their overview networks (Figures 8S, 9S).

The two efforts overlap in 17 ECOD X-groups, and three links between X-groups. The themes in the shared links appear similar. The first link connects X-groups 101 and 819 (fragment #1 in Alva et al.). Both sets include the domain e1nr3A1 (X-group 819), and even the same residues (7-27 in our dataset, 8-27 in theirs). In the X-group 101, both datasets include a domain from the 101.1.4.66 F-group (e1r69A1 in our set vs. e2r1jL1 in theirs). The second link connects X-groups 187 and 205 (fragment #18). There are many cases of domains from the same F-groups: 187.1.1.5 (4 domains in each set) and 205.1.1.8/12/93 (8 domains in ours vs. 6 in theirs). The third connects X-groups 604 and 109 (fragment #28). The domains in both sets are from the F-group 604.12. 1.4, and in the other X-groups, the domains only have the same ECOD T-group 109.4.1. As for the X-groups that although found in both efforts are linked to different X-groups: the domains are not necessarily similar. For example: in the 2003 X-group (Rossmann-like), Alva et al.’s set includes domains from the T-groups 2003.1.[1,2,3,8,9,10,5,14] (fragment #8), while all domains in our set are from 2003.1.1; in the 1 X-group (cradle loop barrels) only the X-group is the same: our domain is from 1.2.1.17, while theirs (fragment #15) are from a different group: 1.1.5.147.

Cases found in the Alva et al. set, and not in ours, are due to different reasons. Some are trivially missing from our set because we use ECOD which is less conservative than SCOP (the domains in fragments (2, 6-7, 9, 11-14, 16, 19, 21-22, 26-27, 33-34, 37) have the same ECOD X-group classification), or because of our more conservative thresholds: fragments (4, 15, 17-18, 20, 23, 25-26, 37, 40) are shorter than 20 residues. In fragments (3, 8, 28-32, 35-36, 38-39) our pre-curated set of bait-themes does not include themes that are from the domains of one of the folds for that fragment (implying that we cannot find cross X-group similarities for this case). That there is no match to any segment in specific domains to any of our bait themes, may be due to these domains removed from our 70% NR set of ECOD domains. Hence, if a particular domain is not in our ECOD set of domains, we checked if any of its close homologues (namely, all the domains with the same ECOD A.X.H.T.F classification) was matched to a bait theme. Finally, the domains in fragments (8, 10, 24), while matched to some bait themes, with sufficiently low E-values, it was not the same bait themes. In summary, although both approaches are based on the HHSearch engine, the significant methodological differences lead to different results.

## Data Availability

The data underlying this article are available in the article, [online](https://trachel-srv.cs.haifa.ac.il/rachel/agilethemes/overview.html), and for [download](https://trachel-srv.cs.haifa.ac.il/rachel/Kolodny_Nepomnyachiy_Tawfik_BenTal_Supp.zip). The downloadable zip file includes Cytoscape [55] /CyToStruct [55] sessions for the different networks and the data files.

## Acknowledgements

We are grateful to Profs. Andrei Lupas and Birte Höcker for the many insightful discussions on these topics, and to Dan Latovicz for his help with the graphical design of all figures. This research has been supported by Grant 94747 by VW foundation. N.B.-T.’s research is supported in part by the Abraham E. Kazan Chair in Structural Biology, Tel Aviv University.

## Footnotes

1 https://trachel-srv.cs.haifa.ac.il/rachel/bridRinRthemes/overview.html

