## Supplemental Data for "Bridging themes: short protein segments found in different architectures"

### Supplementary Material

| X-group id | #dataset domains in X-group | X-group name | Example domain | Architecture | #dataset domains In architecture |
|---|---|---|---|---|---|
| 101 | 24 | HTH | e1efaA1 | α-arrays | 46 |
| 103 | 4 | RuvA-C | e2dhyA1 | | |
| 108 | 3 | EF-hand | e2omzA4 | | |
| 138 | 2 | NO_X_NAME | e3bgeA1 | | |
| 148 | 1 | Histone-like | e4xguA2 | | |
| 187 | 5 | alpha-helical ferredoxin-like | e1e7pB1 | | |
| 3206 | 1 | NO_X_NAME | e2kr0A2 | | |
| 531 | 5 | Domains of FliG | e1lkvX1 | | |
| 589 | 1 | NO_X_NAME | e3rfwA2 | | |
| 150 | 4 | Ferritin/Heme oxygenase/4-helical cytokines | e2cwlA1 | α-bundles | 15 |
| 3403 | 1 | NO_X_NAME | e4dgwA1 | | |
| 3763 | 1 | NO_X_NAME | e3ezuA2 | | |
| 3847 | 1 | NO_X_NAME | e5c0sA1 | | |
| 593 | 2 | NO_X_NAME | e1iokA1 | | |
| 601 | 1 | Four-helical up-and-down bundle | e1ucuA4 | | |
| 604 | 2 | Spectrin repeat-like | e2v6yA1 | | |
| 632 | 3 | immunoglobulin/albumin-binding domain-like | e1l6xB1 | | |
| 3601 | 1 | NO_X_NAME | e2ev4A1 | α-complex topology | 1 |
| 109 | 63 | Repetitive alpha hairpins | e1awcB1 | α-superhelices | 65 |
| 3238 | 2 | Mitochondrial mTERF-like | e2ypfA1 | | |
| 1 | 1 | cradle loop barrel | e1ihmA2 | β-barrels | 19 |
| 2 | 2 | OB-fold | e2wacA2 | | |
| 236 | 4 | GroES-like | e2dphA1 | | |
| 4 | 6 | SH3 | e2diqA1 | | |
| 6120 | 2 | NO_X_NAME | e3zfoA1 | | |
| 70 | 4 | beta-clip | e3n71A1 | | |
| 3702 | 4 | NO_X_NAME | e1rp5A6 | β-complex topology | 4 |
| 207 | 5 | Single-stranded right-handed beta-helix | e1dceA3 | β-duplicates or obligate multimers | 10 |
| 208 | 1 | Single-stranded left-handed beta-helix | e3pr7B1 | | |
| 3512 | 4 | NO_X_NAME | e1p9hA3 | | |
| 10 | 5 | jelly-roll | e3eqeA1 | β-sandwiches | 10 |
| 11 | 1 | Immunoglobulin-like beta-sandwich | e4r5oA2 | | |
| 3414 | 1 | NO_X_NAME | e4l3aB3 | | |
| 3856 | 2 | NO_X_NAME | e4oj6C1 | | |
| 3857 | 1 | NO_X_NAME | e1lktA1 | | |
| 2002 | 3 | TIM beta/alpha-barrel | e1tqjA1 | α/β barrels | 3 |
| 2003 | 64 | Rossmann-like | e1ek6A1 | α/β three-layered sandwiches | 179 |
| 2004 | 36 | P-loop domains-like | e1gkyA1 | | |
| 2007 | 69 | Flavodoxin-like | e1a04A2 | | |
| 2487 | 5 | The swivelling" beta/beta/alpha domains" | e1a6dB3 | | |
| 7523 | 4 | NO_X_NAME | e1eljA2 | | |
| 7577 | 1 | NO_X_NAME | e4nogB1 | | |
| 206 | 1 | NO_X_NAME | e4redA1 | α+β complex topology | 5 |
| 219 | 1 | Cysteine proteinases-like | e3s0qA1 | | |
| 4064 | 1 | NO_X_NAME | e1ej6C1 | | |
| 5093 | 2 | NO_X_NAME | e3rkiA1 | | |
| 3567 | 6 | NO_X_NAME | e3k9aA1 | α+β duplicates or obligate multimers | 6 |
| 223 | 1 | Profilin-like | e3kyqA1 | α+β three layers | 15 |
| 2485 | 13 | Thioredoxin-like | e1z9hA2 | | |
| 4031 | 1 | NO_X_NAME | e3a5xA4 | | |

| | | | | | |
|---|---|---|---|---|---|
| 205 | 16 | NO_X_NAME | e1dwlA1 | α+β two layers | 62 |
| 214 | 1 | NO_X_NAME | e2cr4A1 | | |
| 221 | 24 | beta-Grasp | e1p0rA1 | | |
| 225 | 1 | ATPase domain of HSP90 chaperone/DNA topoisomerase II/histidine kinase-like | e2q2eB1 | | |
| 284 | 2 | FKBP-like | e2jzvA1 | | |
| 304 | 8 | Alpha-beta plaits | e1b7fB2 | | |
| 305 | 1 | DCoH-like | e2pmzS1 | | |
| 308 | 1 | ClpS-like | e2z2lD1 | | |
| 327 | 2 | Alpha-lytic protease prodomain-like | e1yj7A2 | | |
| 3953 | 1 | NO_X_NAME | e3sluA3 | | |
| 4305 | 1 | NO_X_NAME | e2d9bA1 | | |
| 6043 | 1 | yfeY-like | e4ygtA2 | | |
| 809 | 2 | BLIP-like | e2km7A1 | | |
| 819 | 1 | NO_X_NAME | e1nr3A1 | | |
| 2498 | 1 | Zincin-like | e3j1bA2 | mixed α+β and α/β | 1 |
| 3100 | 3 | NO_X_NAME | e2kogA1 | Extended segments | 7 |
| 3362 | 1 | NO_X_NAME | e2b9bA3 | | |
| 3462 | 1 | NO_X_NAME | e3v6iX2 | | |
| 3464 | 2 | NO_X_NAME | e3j0jL2 | | |
| 375 | 6 | Rubredoxin-like | e1k83L1 | few secondary structure elements | 79 |
| 376 | 2 | RING/U-box-like | e2ysmA2 | | |
| 377 | 7 | Glucocorticoid receptor-like | e1b8tA3 | | |
| 386 | 55 | beta-beta-alpha zinc fingers | e1meyC1 | | |
| 389 | 1 | EGF-like | e1igrA3 | | |
| 904 | 2 | B-box zinc-binding domain-like | e2didA1 | | |
| 906 | 6 | CCCH zinc finger | e1rgoA2 | | |

**Table 1S:** The list of ECOD X-groups and the number of domains in each of the X-groups for the set of domains in the bridging themes dataset.

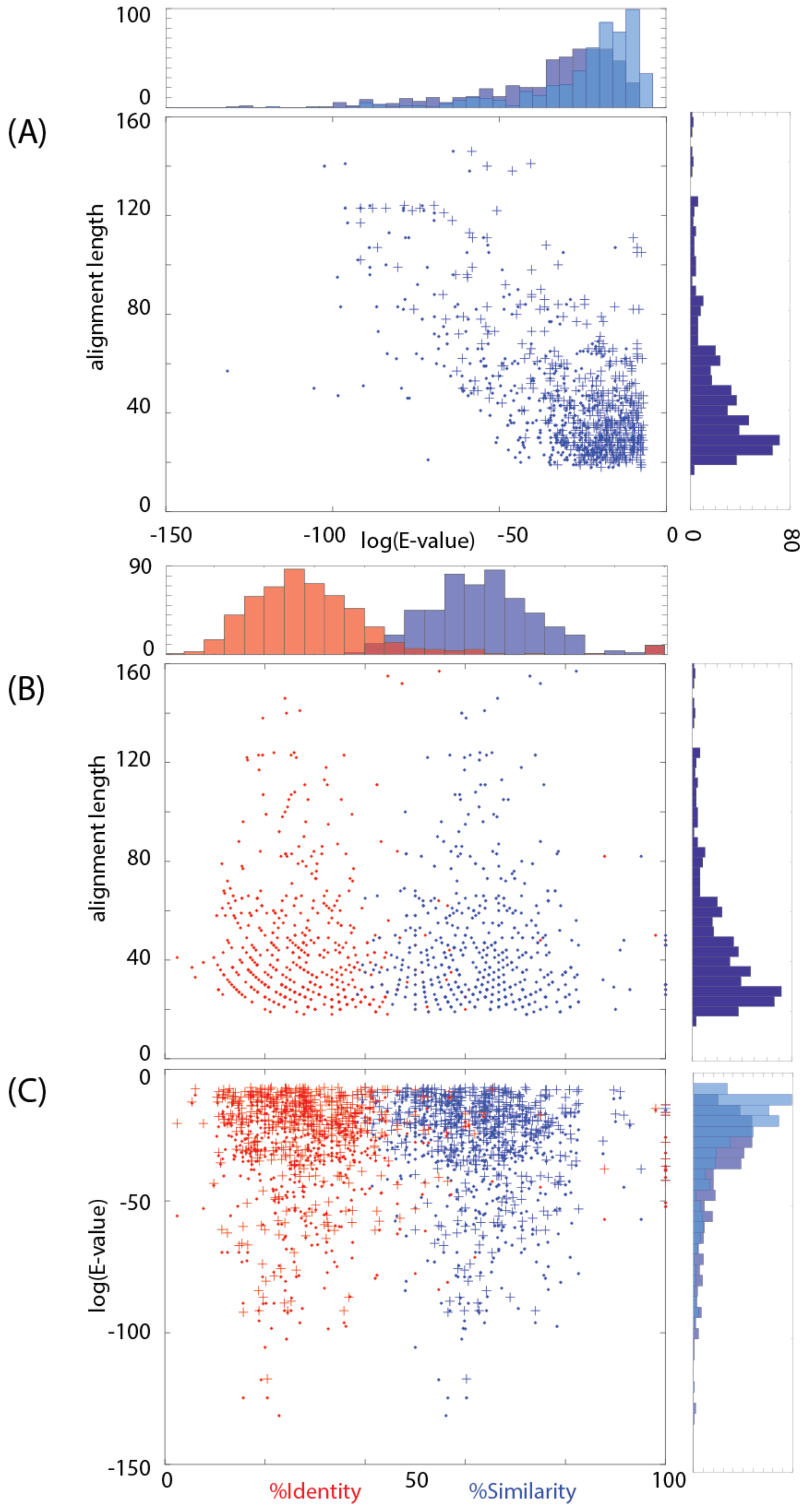


**Figure 1S**: Relating similarity measures:
(A) For the 525 pairs in our dataset, we plot the log(E-value) vs. the alignment length. For each pair of variations, there is a single alignment length value, but two E-values, reported by HHSearch for the “bait” theme and the two domains. We plot these with a ‘+’ (maximal) and ‘.’ (minimal) mark. The histograms of these values are shown to the right and on the top. As expected, longer alignments have lower E-values. (B) The sequence similarity/identify of alignments vs. the alignment length. (C) The sequence similarity/identity of alignments vs. the E-values. The similarity/identify are not correlated with the alignment lengths /E-values.

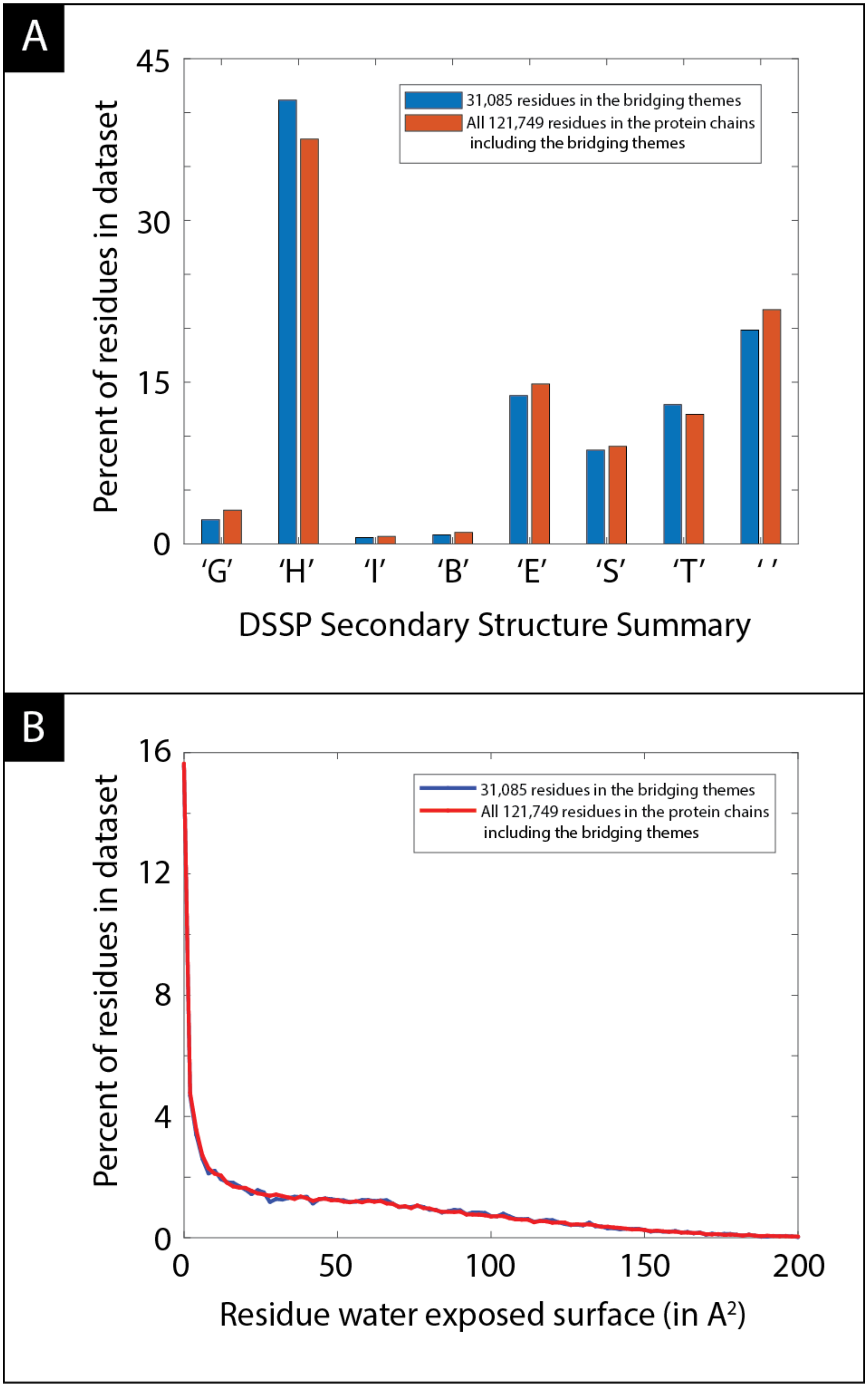


**Figure 2S**: The secondary structure assignments (panel A) and solvent accessibility (panel B) distributions of the residues in the bridging themes (blue) and all residues (red) are similar. The nomenclature and values are derived from DSSP (Kabsch & Sander, 83): G is $3_{10}$-helix, H is α-helix, I is π-helix, B residue in isolated β-bridge, E is extended strand, S is bend, T is hydrogen bonded turn and a blank (the last column) is a loop or irregular.

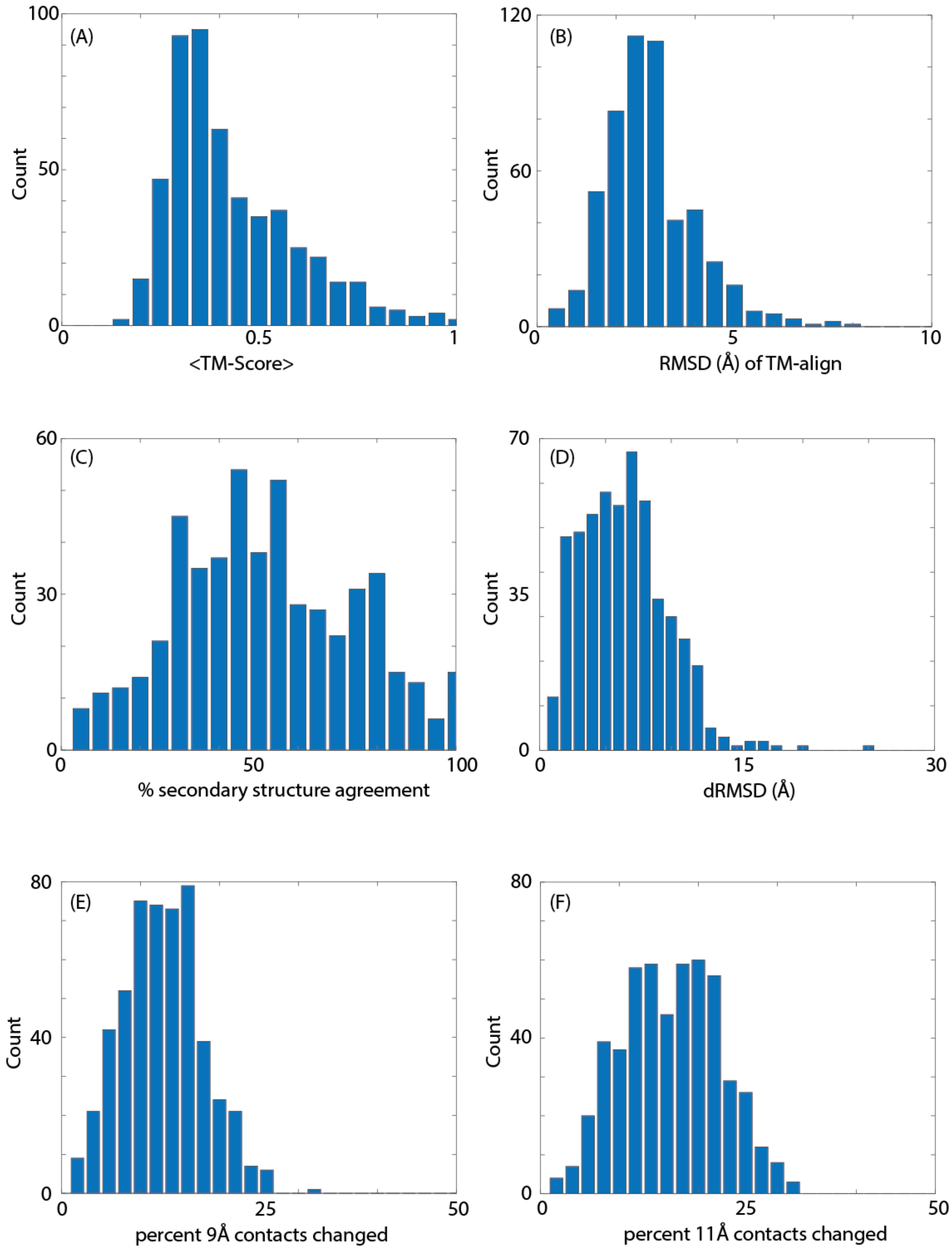


**Figure 3S**: Histograms of structural similarity measures comparing the variations of the bridging themes. The measures in Panel A,B are calculated by TM-align. TM-scores range between 0-1, with scores > 0.5 indicating the same fold (25% of the pairs), and <0.3 indicating random structural similarity (30% of the pairs). The RMSDs of the structurally aligned residues (panel B) are lower than those the sequence aligned residues (Figure 3 in the main document) because these are only a subset of the residues (and the ones that are structurally similar). Panel C shows the percent secondary structure agreement, and we see that in many cases, this agreement is not high. Panel D shows the dRMSD, Panels E,F show the percent contacts changed using two thresholds (9 Å and 11 Å).

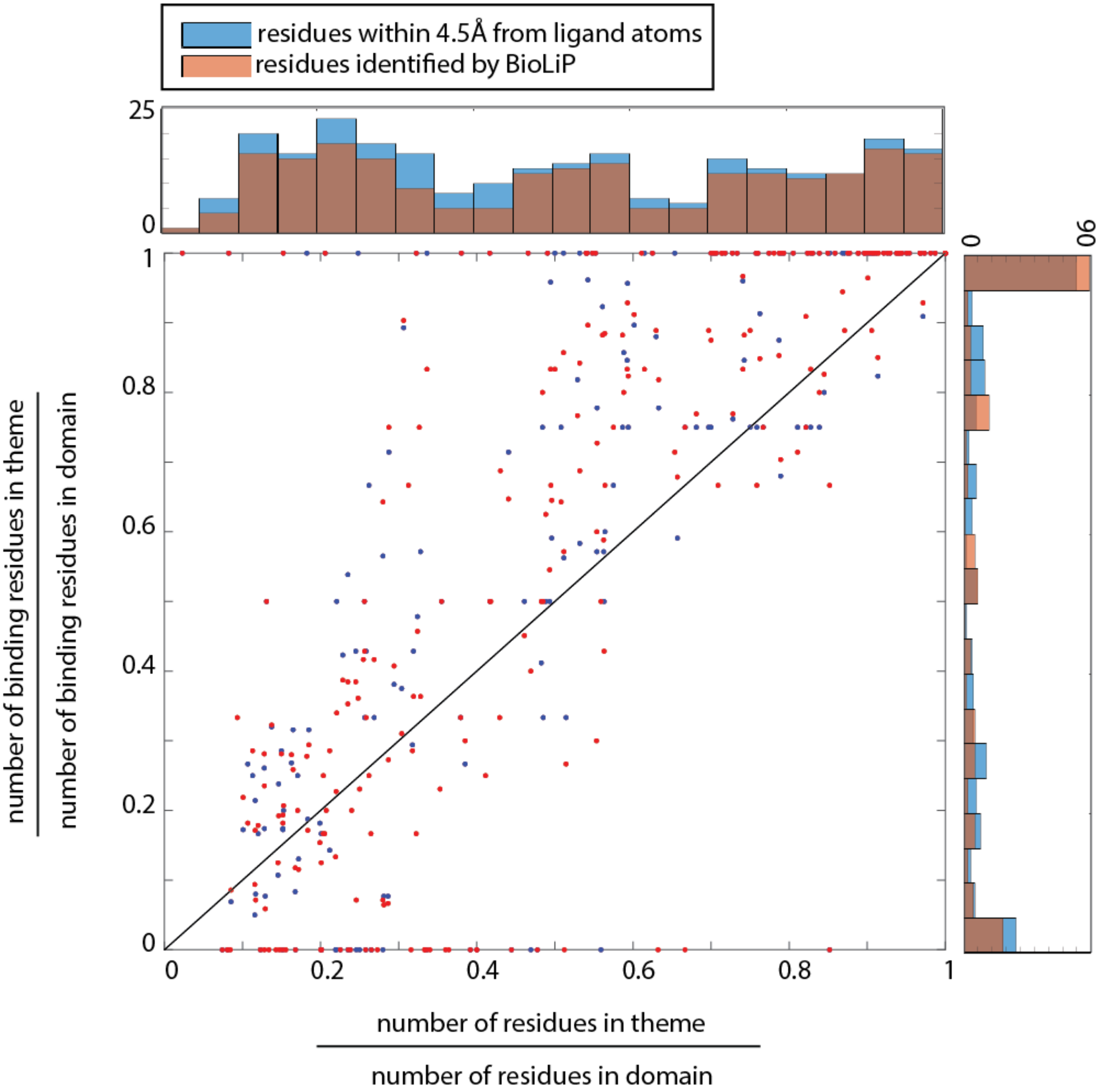


**Figure 4S:** Comparing two rations: along the x-axis, the ratio of the number of residues in theme to number of residues in domain vs. along the y-axis the ratio of the number of binding residues in theme to total number of binding residues in domain. T cumulative histograms of these ratios are also shown. The ratio along the x-axis serves as the background distribution of the percent of residues that are in a bridging theme for domains for which we have binding data: In blue we show the histogram for the 263 domains in which we identified binding of a ligand with 4.5 Å of one of its residues, and in red-brown, the dataset of 217 domains for which we have BioLiP data. In both datasets, we see an enrichment of the high ratios for binding residues, i.e., there are more cases in which binding residues are within a bridging theme (shown as datapoints to the upper-left of the diagonal line).

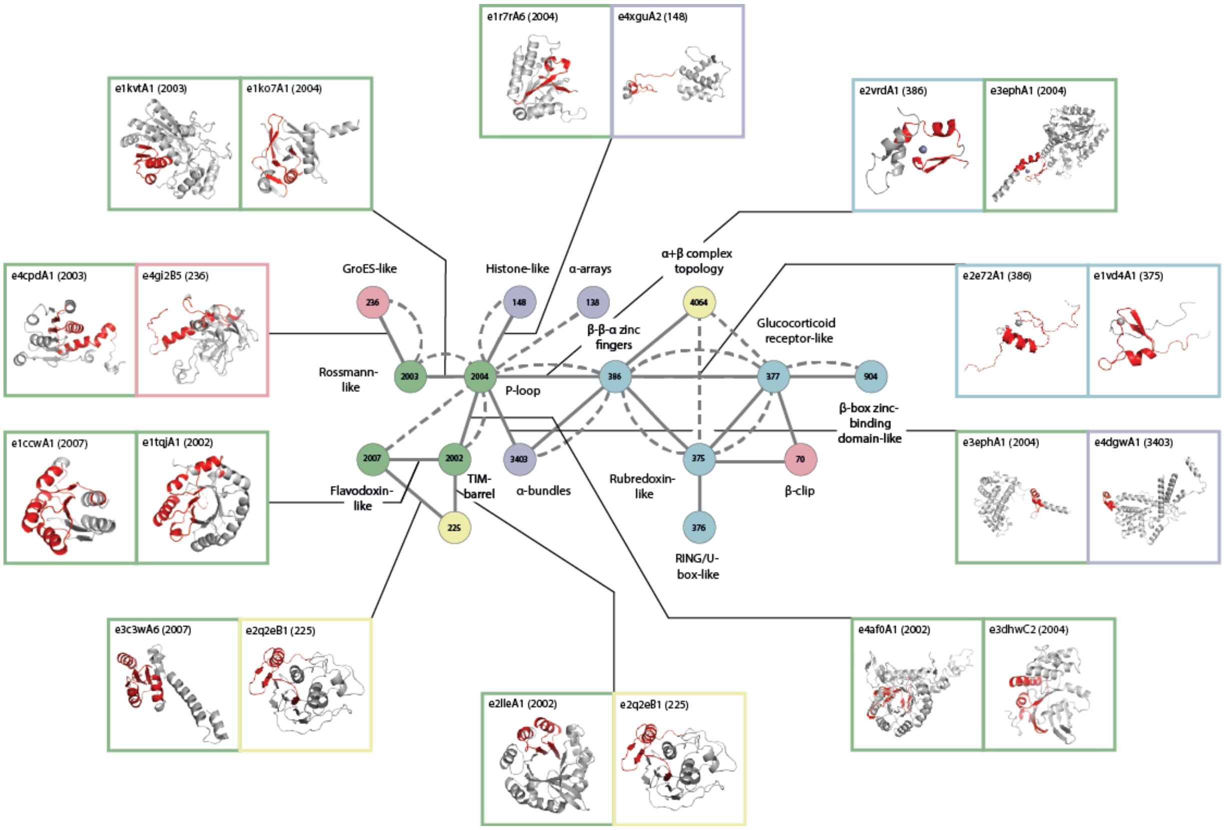


**Figure 5S:** A part of the largest connected component in the overview networks (Figure 4). An arbitrarily chosen example is shown for each pair of X-groups that share a theme. Shown are the structures of the two domains, colored in gray, and the shared theme highlighted in red. Many of the ECOD X-groups in this connected component are considered ancient, including the P-loops, the Rossmanns, and the TIM-barrels.

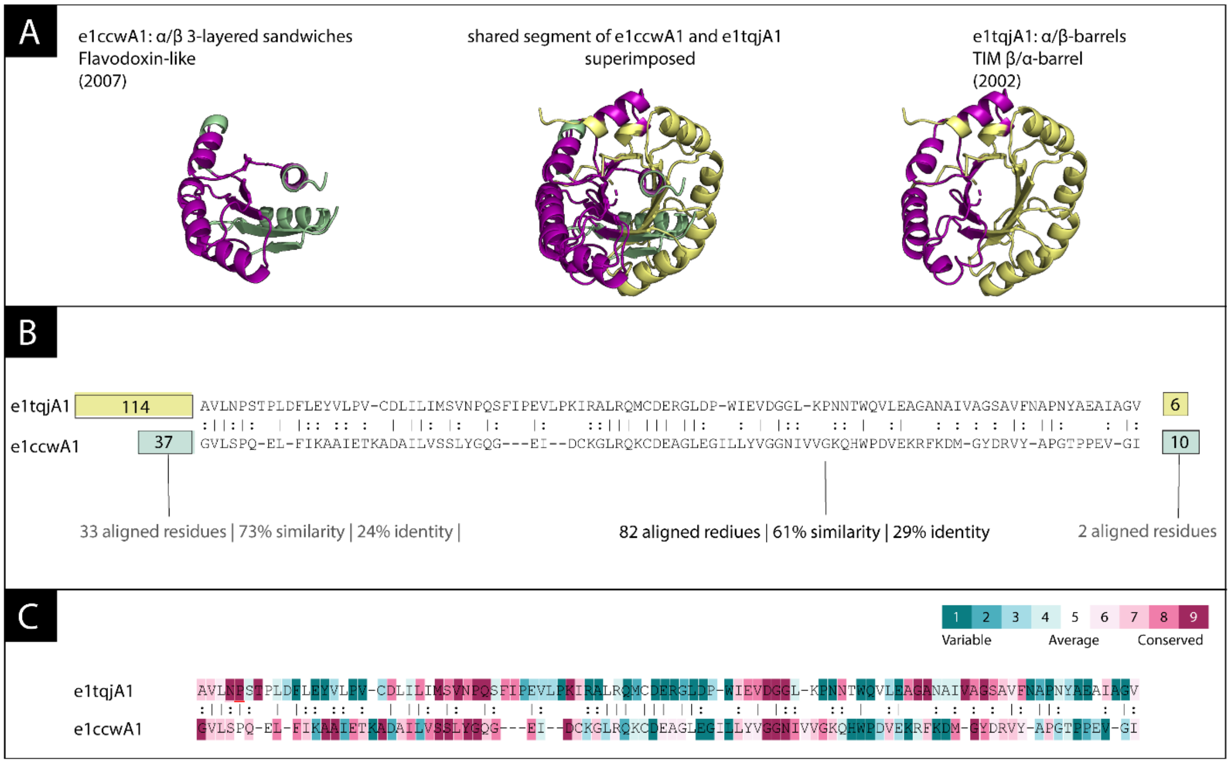


**Figure 6S:** A bridging theme shared between e1ccwA1 (Flavodoxin-like domain) and e1tqjA1 (TIM beta/alpha barrel) (A) The overall structures are characteristic of their folds: the structures are shown superimposed in the middle and separated for clarity. The shared theme is colored in violet. The structures of the variations are very similar (Cα RMSD = 1.38 Å) while the rest of the structures is not. (B) Sequence alignment shows that the similarity between the variations on the shared theme are similar to the respective N-terminal regions of the domains (C) ConSurf analysis of the corresponding proteins (1tqj and 1ccw) and their respective homologues.

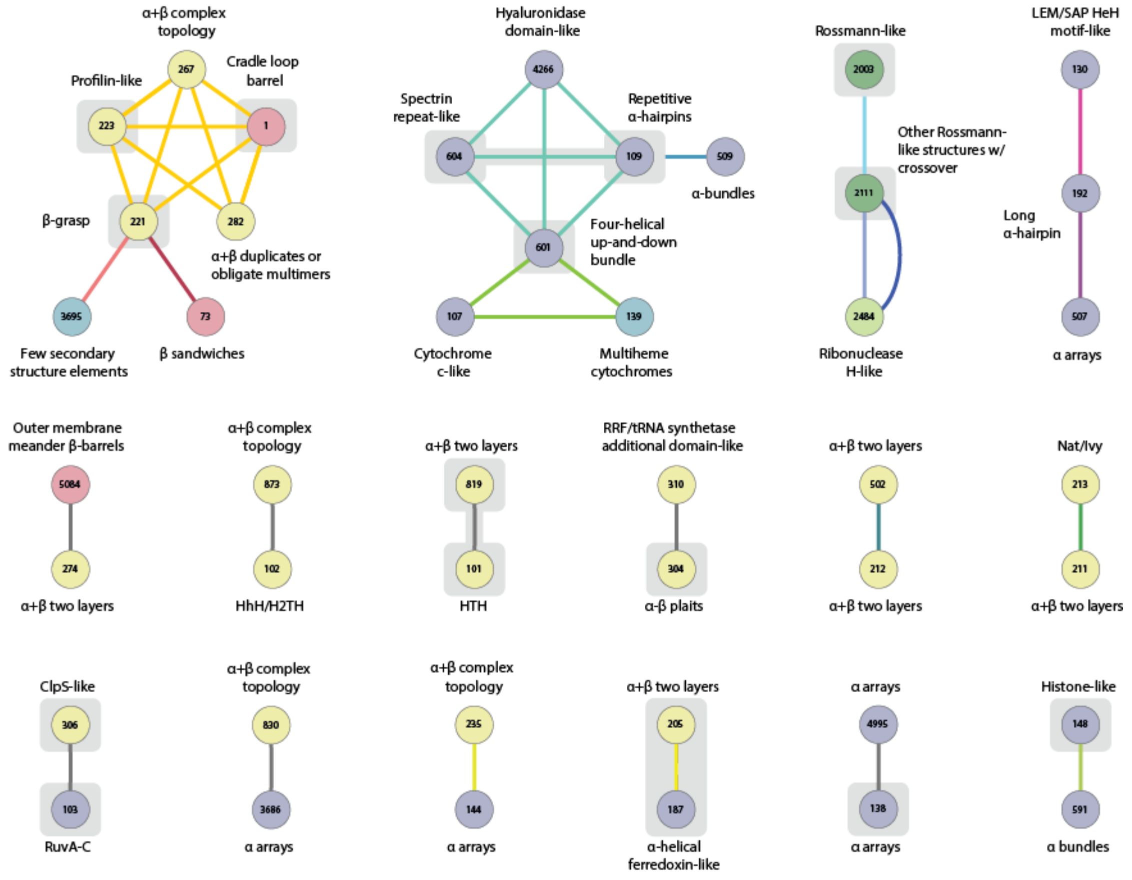


**Figure 7S**: An overview network for the Alva et. al. set [1]. We use Figure 4's color coding, and add the yellow-green shade for the mixed a+b and a/b class (2484). The color of the edge connecting two X-groups encodes the serial number of the fragment in the set: e.g., domains from X-groups 601, 107, and 139 appear in the same fragment (#17). The seventeen ECOD X-groups and three edges that were also found among our agile themes are shown with a gray background.

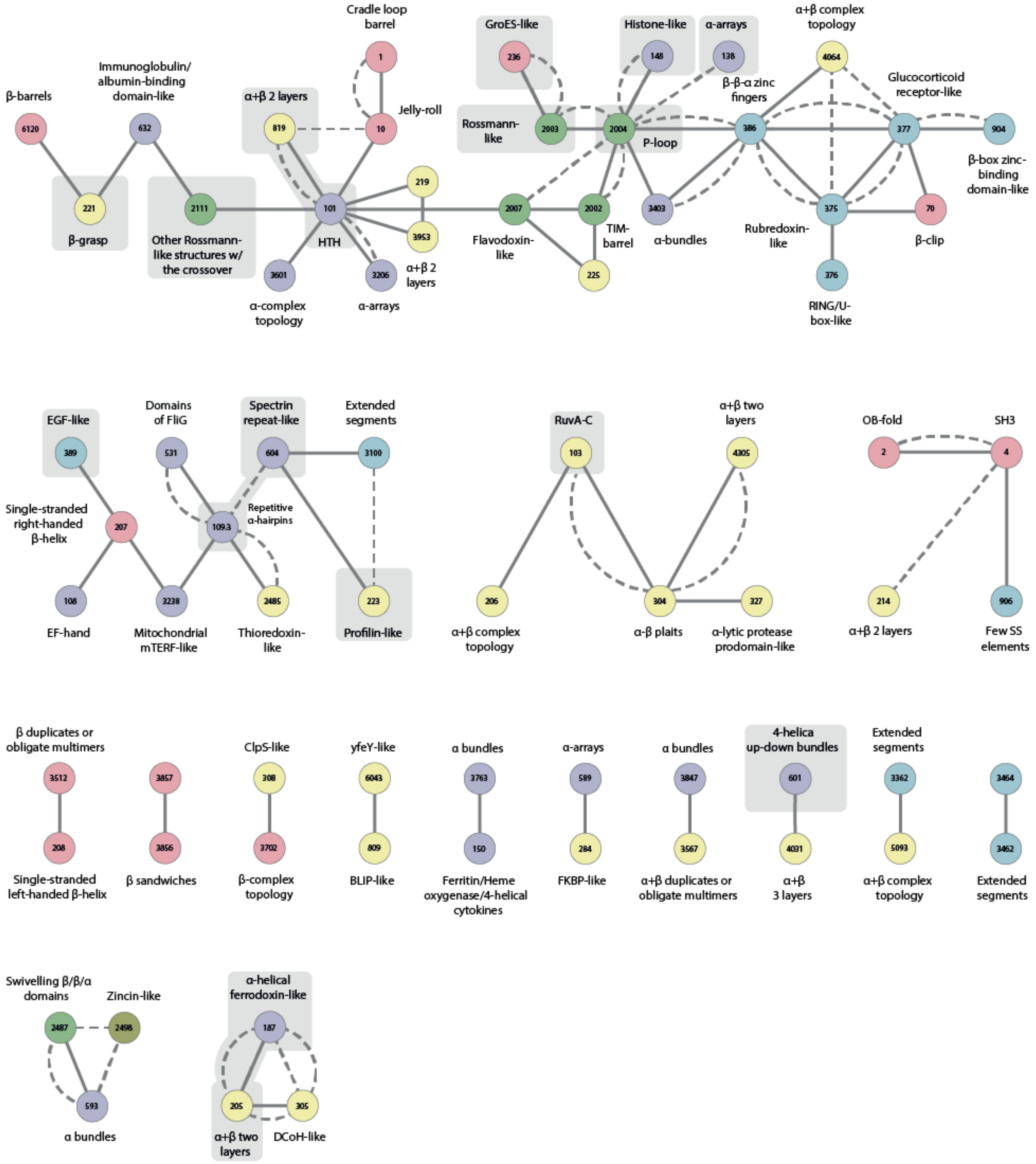


**Figure 8S:** Highlighting the regions in protein space with shared themes found by us and by Alva et al., within our overview network (see Fig. 4). There are 17 ECOD X-groups and 3 edges that were also found in Alva et al.'s dataset shown with a gray background. Node colors correspond to class of the X-group: blue for all-α, red for all-β, green for α/β, yellow for α+β, and cyan for others, and edges connect pairs of different X-groups for which we found a common theme in a domain in each of the two groups.

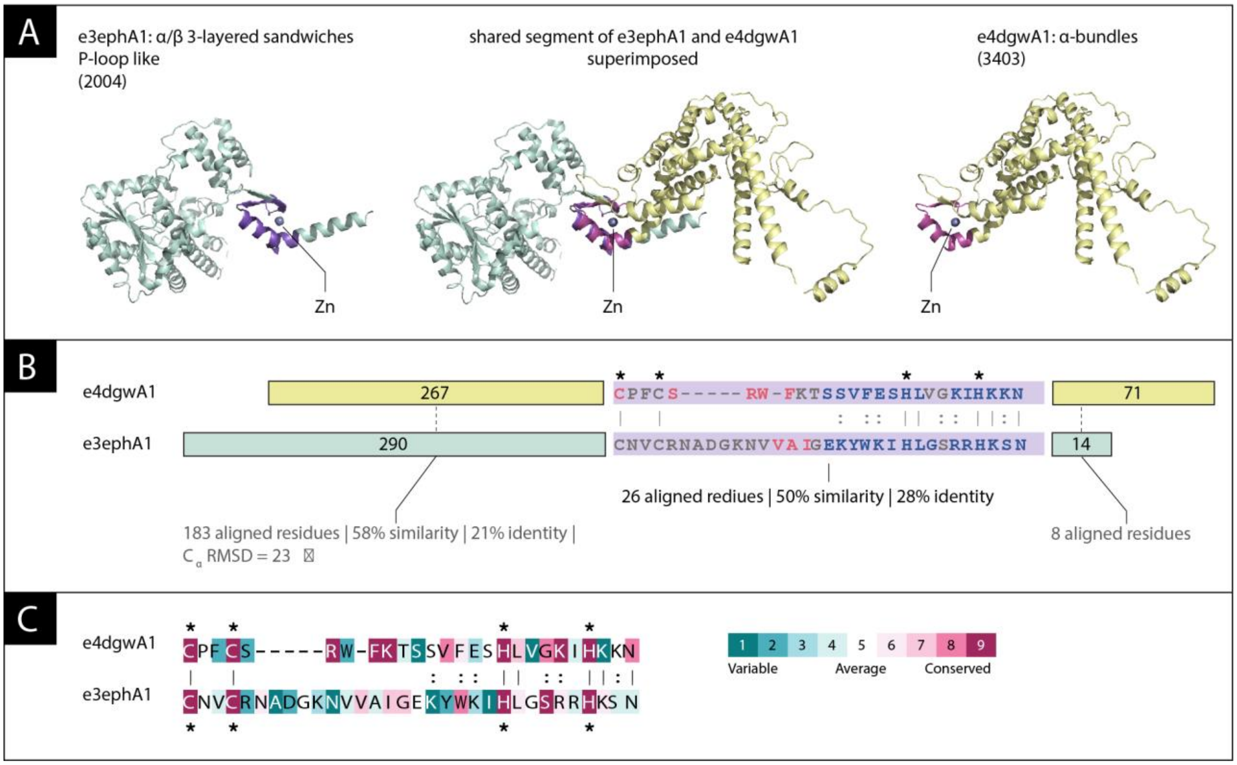


**Figure 9S:** A zinc-binding theme shared between the domain e3ephA1 from the 2004 X-group and the domain e4dgwA1 from the 3404 X-group. (A) The overall structures are different. Domain e3ephA1 (left) is a P-loop like domain from the alpha/beta A-group, and e4dgwA1 (right) is from the all-alpha domain-group. The 26 residues long shared theme (pink) binds zinc (grey sphere). When structurally superimposing the shared theme (middle), the structures of the two are very similar (Cα RMSD = 1.38 Å) while the rest of the structures is not. (B) Sequence alignment shows that the similarity between the variations on the shared theme are similar to the respective N-terminal regions of the domains, and much higher than the similarity of the respective C-terminal regions. The four residues that mediate zinc binding in both domains, marked with asterisks, align to each other in both variations. (C) ConSurf analysis of the corresponding proteins (4dgw and 3eph) and their respective homologues show that the zinc binding residues (marked with asterisks) are highly conserved.

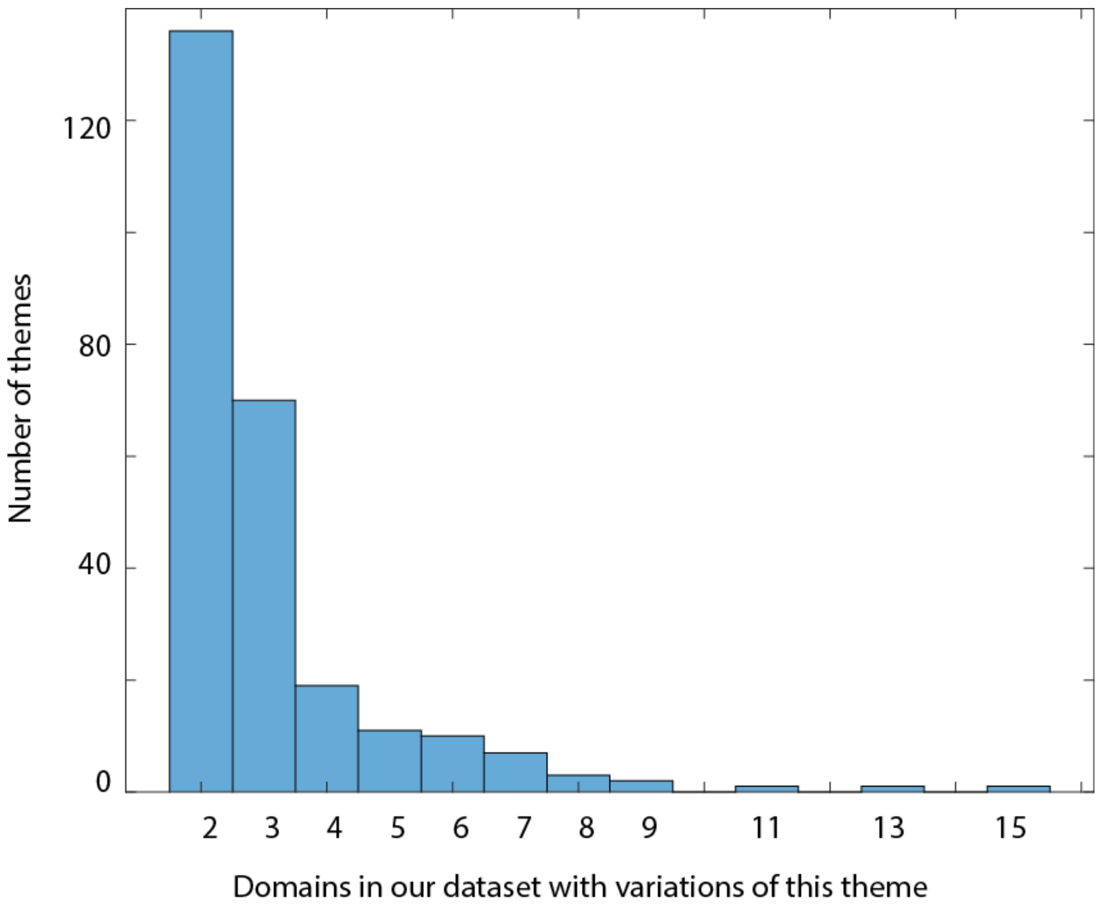


**Figure 10S:** The number of domains identified by each of the bait themes that contributed variations to our bridging data set. Altogether, 261 bait themes successfully identified bridging themes, and the number of domains that each identified varies: over half (136) identified only the two domains in different X-groups. The maximal number of domains related in our set by one bait theme is 15.
